# Hepatic estrogen receptor α is required for stage-specific coupling of liver metabolism and proliferation during pregnancy

**DOI:** 10.64898/2026.08.10.743939

**Authors:** Clara Meda, Arianna Dolce, Giacomo Talamazzini, Claes Ohlsson, Fabrizia Carli, Patrizia Infelise, Amalia Gastaldelli, Adriana Maggi, Sara Della Torre

## Abstract

**Background and Aims:** Pregnancy requires dynamic, stage-specific adaptations in maternal liver metabolism and growth to sustain fetal development while preserving systemic homeostasis. Estrogen signaling, which significantly increases during pregnancy, is primarily mediated in hepatocytes by estrogen receptor α (ERα). Although hepatic ERα regulates female liver metabolism under non-pregnant conditions, its role in pregnancy-induced hepatic remodeling remains unclear.

**Methods:** We studied non-pregnant and pregnant control and liver-specific ERα knockout (LERKO) mice across gestational stages using longitudinal physiological measurements, liver transcriptomics, targeted metabolomics, histological assessment of cell proliferation, and metabolic phenotyping.

**Results:** In control mice, pregnancy elicited sequential hepatic remodeling characterized by early induction of cell-cycle programs, a mid-gestational peak in hepatocyte proliferation with transient suppression of selected metabolic pathways, and late reactivation of specific metabolic programs. Chronic hepatic ERα deficiency alters this temporal pattern. LERKO livers showed premature activation of proliferative and anabolic transcriptional programs, changes in amino acid- and fatty acid-related metabolic pathways, and altered temporal regulation of AKT-mTORC1-related signaling. At mid-gestation, LERKO mice displayed reduced hepatocyte proliferation, altered expression of metabolic and insulin-related genes, blunted gestational glucose adaptation without overt evidence of systemic insulin resistance, and changes in the light/dark-phase metabolic patterns.

**Conclusions:** These findings suggest that hepatic ERα is required for the appropriate stage-specific coupling of liver growth, metabolic remodeling, and insulin-responsive signaling during pregnancy. Its loss is associated with gestational hepatic maladaptation and systemic metabolic phenotypes, providing a framework for investigating estrogen-dependent mechanisms underlying pregnancy-associated metabolic and liver disorders.

**Highlights:** Hepatic ERα is required for stage-specific liver remodeling during pregnancy. Loss of hepatic ERα alters temporal coupling of liver growth and metabolism. LERKO mice show early changes in amino acid- and fatty acid-related pathways. Hepatic ERα loss reduces proliferation and alters gestational glucose adaptation.

Hepatic ERα loss is associated with altered light/dark-phase metabolic organization.

**Graphical abstract:** 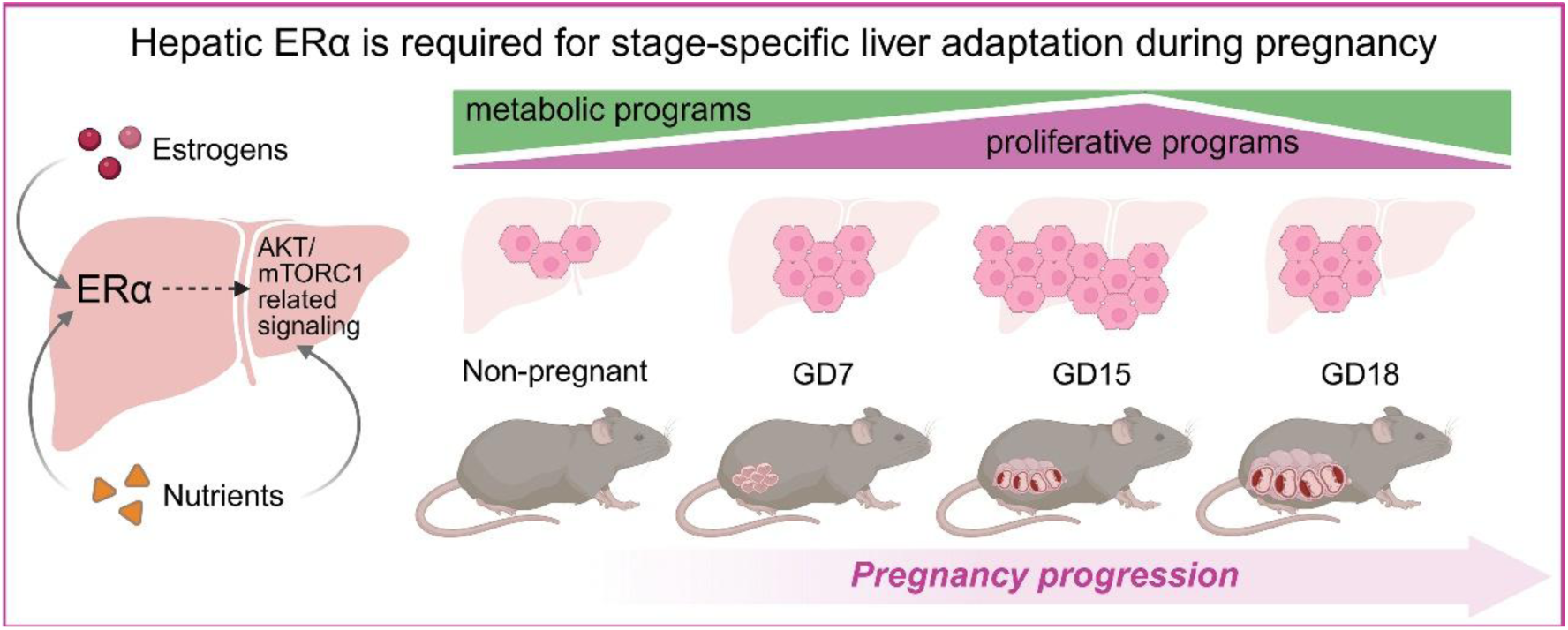

## Introduction

Pregnancy induces significant metabolic adaptations across maternal organs to support fetal growth while maintaining maternal systemic homeostasis [1,2]. Among these organs, the liver plays a pivotal role by integrating endocrine and nutritional signals to regulate glucose, lipid, amino acid (AA), and bile acid metabolism [2–5]. Alongside these functions, the maternal liver undergoes marked structural and functional remodeling, including hepatocyte hypertrophy and proliferation, which contributes to the progressive increase in liver mass during gestation [6,7]. These adaptations are dynamic and occur in a stage-specific manner, suggesting that liver growth and metabolic rewiring must be temporally coordinated to meet the evolving demands of both the mother and the festus [2].

Estrogen levels markedly rise during pregnancy [8], influencing hepatic metabolism, insulin sensitivity, mitochondrial function, and lipid handling [2,3,9]. Estrogen actions in hepatocytes are largely mediated by estrogen receptor α (ERα), the predominant estrogen receptor expressed in hepatocytes [10,11]. In non-pregnant conditions, hepatic ERα promotes metabolic flexibility, influences sexually dimorphic gene expression, and protects against metabolic dysfunctions, including metabolic dysfunction-associated steatotic liver disease (MASLD) [10–13]. Previous studies using liver-specific ERα knockout models have established hepatic ERα as a regulator of female liver metabolic homeostasis outside pregnancy [13–15]. However, the metabolic actions of hepatic ERα are highly context-dependent [16], and pregnancy represents a distinct physiological state characterized by elevated estrogen levels, physiological insulin resistance, altered substrate utilization, and increased fetal nutrient demand [9,17–19]. Preclinical studies indicate that ERα expression and activity are dynamically regulated during gestation, suggesting potential stage-dependent modulation of estrogen-responsive pathways [20,21]. At the same time, pregnancy places substantial metabolic stress on the maternal liver, and failure to adapt appropriately may contribute to gestational metabolic disorders, including gestational diabetes mellitus (GDM), MASLD, and intrahepatic cholestasis of pregnancy (ICP) [22–27]. Although these human disorders are multifactorial, they underscore the vulnerability of hepatic metabolic adaptation during pregnancy and suggest that impaired hepatic estrogen signaling may contribute to maladaptive maternal metabolism in susceptible contexts.

Despite the established role of hepatic ERα in non-pregnant female liver physiology [10,11,13–15], its requirement for stage-specific coupling of liver growth and metabolic remodeling during pregnancy remains unclear. It is not yet determined whether a deficiency or alteration in hepatic ERα affects the normal temporal coordination among hepatocyte proliferation, regulation of metabolic pathways, insulin-responsive signaling, and systemic metabolic adaptation during gestation. This distinction is significant because pregnancy may uncover hepatic ERα functions that are not evident in non-pregnant models.

To address this issue, we used a liver-specific ERα knockout (LERKO) mouse model and analyzed non-pregnant and pregnant females across various gestational stages using longitudinal physiological measurements, liver transcriptomics, targeted metabolomics, histological assessment of hepatocyte proliferation, AKT–mTORC1-related signaling analyses, glucose and insulin tolerance tests, and indirect calorimetry.

Our findings indicate that hepatic ERα is required for the proper stage-specific integration of liver cell proliferation with metabolic rewiring, insulin-responsive signaling, and systemic metabolic adaptation during pregnancy. Chronic deficiency of hepatic ERα is associated with premature activation of proliferative and anabolic transcriptional programs, alterations in AA and fatty acid-related metabolic pathways, impaired mid-gestational hepatocyte proliferation, and changes in light/dark-phase metabolic patterns. These results identify hepatic ERα as a critical factor in maternal liver adaptation to pregnancy and provide a framework for exploring how altered hepatic estrogen signaling may contribute to gestational metabolic maladaptation.

## Materials and methods

### Animals and experimental design

Syngeneic ERα-floxed control (CTRL) and LERKO female mice were both C57BL/6J strain [28]. Four- to five-month-old mice were fed *ad libitum* with a standard diet (Ssniff V1534-703), provided with filtered water, and maintained at 22–25°C, 50±10% relative humidity, under an automatic 12-h light/dark cycle. Females were mated during *proestrus/estrus*; gestational day (GD) 0 was defined as the day of mating. Pregnant females were euthanized at GD7, GD15, and GD18, whereas non-pregnant (NP) females were collected during *proestrus* to provide a hormonally standardized baseline comparison. Mice were euthanized after 6-h of fasting in the early afternoon to minimize feeding- and time-dependent metabolic variability [28]. For acute insulin stimulation, 6-h-fasted mice received intraperitoneal human insulin (0.75 U/kg) or vehicle, and the livers were collected 12 min later. All protocols were approved by the Istituto Superiore di Sanità ethics committee (1272/2015-PR and 149/2022-PR) and followed the ARRIVE guidelines and European regulations.

### Sex hormone measurements

17β-estradiol, estrone, progesterone, androstenedione, testosterone, and DHT were quantified using validated gas chromatography-tandem mass spectrometry, with quantification limits of 0.5, 0.5, 74, 12, 8, and 2.5 pg/mL, respectively, as previously described [29].

### Biochemical analysis

Triglyceride (TG) levels were measured using commercial kits according to the manufacturer’s protocol (Abcam, ab65336).

### RNA extraction, qPCR, and RNA-sequencing

Total liver RNA was isolated, reverse transcribed, and analyzed by qPCR as previously described [30]. Gene expression was normalized to *Rplp0* and calculated using the 2⁻^ΔΔCt^ method [31]. Primer sequences are listed in Supplementary Table S1. For RNA-seq, libraries were prepared from 500 ng of quality-controlled total RNA using the Illumina Stranded mRNA Prep Ligation protocol and sequenced on an Illumina NovaSeq platform to generate 2 × 150 bp paired-end reads.

### Transcriptomics data analysis

Raw sequencing reads were quality-controlled, adapter-trimmed, and aligned to the mouse reference genome GRCm38/Gencode M20 using STAR. Gene-level counts were generated with STAR GeneCounts and used for differential expression analysis with DESeq2. Quality control included assessment of gene-body coverage, exon-mapping efficiency, sample correlation, and number of detected genes to identify potential outliers or mapping failures. DEGs were defined using FDR-adjusted p-values <0.05. Exploratory analyses included clustering and principal component analysis. Detailed procedures for Gene Ontology, pathway, motif, Venn, and UpSet analyses are provided in the Supplementary Methods.

### Metabolomic analysis

Detailed procedures are provided in the Supplementary Methods. A complete metabolite table, including individual values, group means, fold changes, nominal p-values, and FDR-adjusted p-values, is provided in Supplementary Table S2.

### Immunohistochemical analysis

Formalin-fixed, paraffin-embedded liver sections were analyzed using H&E staining and Ki67 immunohistochemistry. Ki67-positive nuclei were quantified as a percentage of total nuclei using QuPath, together with hepatic morphometric parameters. Detailed procedures are provided in the Supplementary Methods.

### Glucose and Insulin Tolerance Analysis

Glucose (GTT) and insulin (ITT) tolerance tests were performed in NP and GD15 females after overnight (GTT) or 4-h (ITT) fasting. Mice received intraperitoneal injections of glucose (2 g/kg) or insulin (0.75 IU/kg), and blood glucose levels were measured at baseline and 15, 30, 60, 90, and 120 min post-injection using a glucose meter (Accu-Chek® Instant Meter, Roche).

### Western Blotting Analysis

Total and phosphorylated proteins were quantified by western blotting in frozen liver samples. Detailed procedures, antibody information, and uncropped raw blot images are provided in the Supplementary Methods and Data files.

### Metabolic cages and indirect calorimetry

Indirect calorimetry was performed using the Promethion Metabolic Screening system. Mice were singly housed, acclimatized for 48 h, and monitored for 48 h under a 12-h light/dark cycle with ad libitum access to food and to water. VO₂, VCO₂, RER, locomotor activity, food intake, and energy expenditure were recorded and analyzed as hourly averages and light/dark-phase summaries using CalR. Sensitivity analyses comparing 48-, 72-, and 84-h acclimatization periods did not reveal any major differences in the measured parameters. The raw data and analyzed outputs are provided in Supplementary Table S3.

### Statistical analysis

Statistical analyses were performed using GraphPad Prism 8.0. Data are presented as mean ± SEM. Exact group-specific sample sizes are reported in the figure legends and Supplementary Table S4; each biological replicate represents one animal, unless otherwise specified. Two-group comparisons used two-tailed unpaired Student’s t-tests, whereas multiple-group datasets were analyzed by one- or two-way ANOVA, followed by Bonferroni-adjusted comparisons. Factorial designs were analyzed by two-way ANOVA to assess main effects and interactions. Longitudinal GTT/ITT profiles and phase-averaged indirect calorimetry data were analyzed by repeated-measures ANOVA, with time or light/dark phase as the within-subject factor and experimental group as the between-subject factor. All tests were two-sided, with p < 0.05 considered statistically significant. Additional details are provided in the Supplementary Methods.

## Results

### Hepatic ERα deficiency alters estrous cycle dynamics and mid-gestation reproductive outcomes

LERKO females exhibited prolonged estrous cycles compared with CTRL (Fig. 1A), in line with previous findings [28]. To investigate the role of hepatic ERα during pregnancy, females were mated during the *proestrus/estrus* phase and analyzed at GD7, GD15, and GD18; non-pregnant (NP) controls were collected at *proestrus* (Fig. 1B). Pregnancy rates were comparable to those previously reported for the C57BL/6J strain [32], although slightly lower in LERKO mice (32%) than in CTRL mice (36%), without reaching statistical significance (Fig. 1C). The mean litter size did not significantly differ between genotypes; however, litter size variability was greater in LERKO mice (coefficient of variation, CV: 47.8% vs. 26.5% in CTRL mice, F-test p<0.001; Fig. 1D–E). At GD15, LERKO dams exhibited a non-significant trend toward increased fetal resorption (Fig. 1F), along with significantly reduced fetal weights compared with those of CTRL mice (p<0.001; Fig. 1G).

**Fig. 1.**
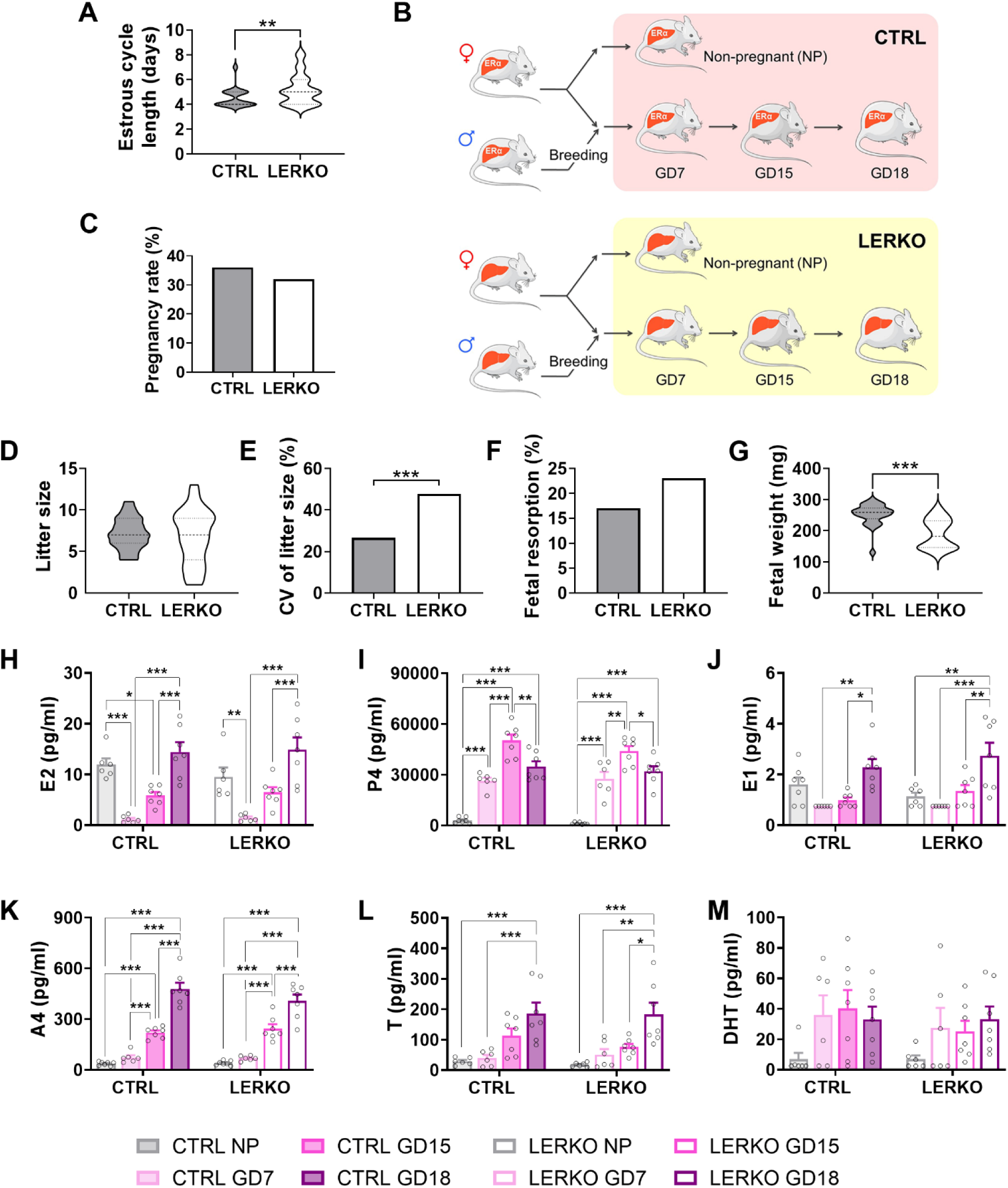
Lack of hepatic ERα alters estrous cycle progression and mid-gestation reproductive outcomes. (**A**) Length of estrous cycle in CTRL (n=36) and LERKO (n=38) female mice, assessed by vaginal smears analysis. (**B**) Experimental design used to assess the role of hepatic estrogen signaling during pregnancy. (**C**) Pregnancy rate of CTRL (n=50) and LERKO (n=62) females. (**D-E**) Litter size (**D**) and its coefficient of variation (CV%; **E**) in CTRL (n=50) and LERKO (n=62). (**F**) Percentage of resorbed fetuses at GD15 in CTRL (n=50) and LERKO (n=62) females. (**G**) Fetal weight measured at GD15. Data are mean ± SEM (n=26 for CTRL; n=28 for LERKO), ***p<0.001 LERKO *vs* CTRL by two-tailed unpaired Student’s *t*-test. (**H-M**) Plasma levels of 17β-estradiol (E2; **H**), progesterone (P4; **I**), estrone (E1; **J**), androstenedione (A4; **K**), testosterone (T; **L**), and dihydrotestosterone (DHT; **M**) in non-pregnant (NP) and pregnant CTRL and LERKO at GD7, GD15 and GD18. Data are mean ± SEM (CTRL NP/GD7/GD15/GD18, n=6/6/7/7; LERKO NP/GD7/GD15/GD18, n=6/6/7/7; each data point represents one independently analyzed pool, and each pool comprised plasma from 2–3 individual animals). *p<0.05, **p<0.01 and ***p<0.001 by two-way ANOVA followed by Bonferroni’s *post hoc* test.

Despite these reproductive alterations, circulating levels of 17β-estradiol (E2), progesterone (P4), estrone (E1), androstenedione (A4), dihydrotestosterone (DHT), and testosterone (T) displayed stage-dependent gestational profiles, with no significant differences between genotypes at the analyzed stages (Fig. 1H–M). Thus, reproductive alterations in LERKO mice occurred without detectable genotype-dependent changes in circulating sex-steroid levels.

### Pregnancy drives liver growth and metabolic remodeling

Throughout pregnancy, both genotypes exhibited a progressive increase in maternal body weight (BW) and liver mass (Fig. 2A–C). Liver weight positively correlated with BW in both genotypes, particularly in CTRL mice (Fig. 2D–E). Liver weight increased during pregnancy, and when normalized to BW, it peaked at GD15, with LERKO mice exhibiting higher liver-to-BW *ratios* than CTRL mice (p=0.014), consistent with altered hepatic growth dynamics associated with hepatic ERα deficiency (Fig. 2F-G).

**Fig. 2.**
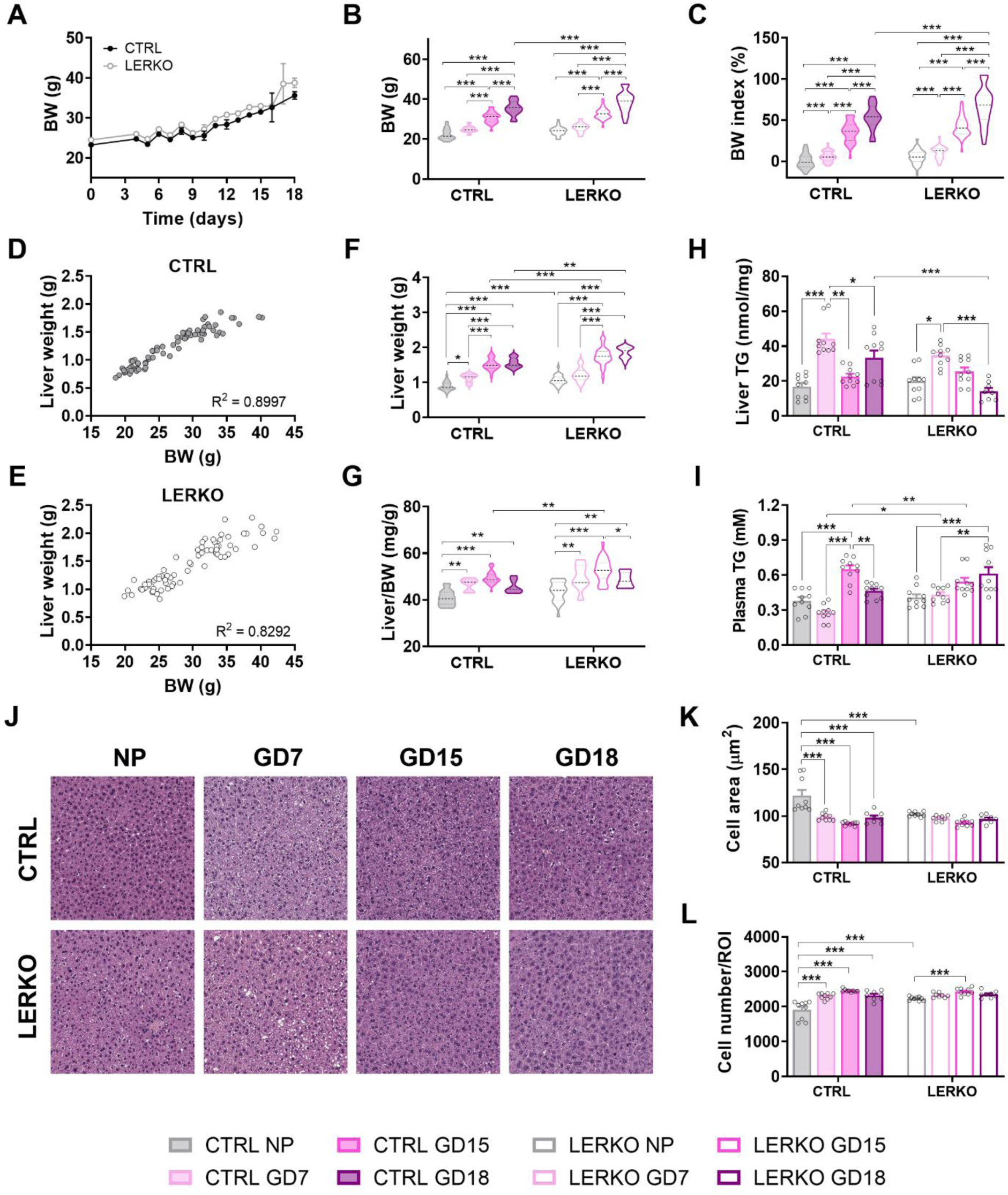
Pregnancy progression is associated with liver growth and metabolic remodeling. (**A-B**) Body weight (BW) of CTRL and LERKO females measured throughout pregnancy. (**C**) BW index (%) of NP and pregnant CTRL and LERKO at GD7, GD15 and GD18. (**D-E**) Pearson’s correlation between body and liver weights across reproductive stages in CTRL (**D**, n=70) and LERKO (**E**, n=70). (**F-G**) Absolute liver weight (**F**) and liver-to-BW *ratio* (**G**) in CTRL and LERKO mice across pregnancy. (**H-I**) Triglyceride (TG) levels measured in the liver (**H**) and plasma (**I**) of CTRL and LERKO during gestation. (**J-L**) Representative hematoxylin and eosin (H&E)-stained liver sections (**J**) and quantitative analyses of cell area (**K**) and cell density (**L**). For panels **A-C** and **F-G**: CTRL NP/GD7/GD15/GD18, n=20/16/18/16, and LERKO NP/GD7/GD15/GD18, n=20/16/18/16. For panel **H**: CTRL NP/GD7/GD15/GD18, n=10/10/10/10, and LERKO NP/GD7/GD15/GD18, n=10/10/10/8. For panel **I**, n=10 per group. For panels **K–L**, CTRL NP/GD7/GD15/GD18, n=9/9/9/8, and LERKO NP/GD7/GD15/GD18, n=9/8/9/8. Data are mean ± SEM; *p<0.05, **p<0.01 and ***p<0.001 by two-way ANOVA followed by Bonferroni’s *post hoc* test. ROI: region of interest.

Hepatic triglyceride (TG) content increased by GD7 in both genotypes, with a more pronounced increase in CTRL mice. TG levels declined by GD15 in CTRL livers, whereas a later decline was observed at GD18 in LERKO livers (Fig. 2H). Plasma TG levels peaked at GD15 in CTRL mice, whereas LERKO mice showed elevated levels at both GD15 and GD18, with the highest mean value at GD18 (Fig. 2I). Histological analysis (H&E) revealed dynamic changes in hepatic cell morphology across pregnancy (Fig. 2J-L). In CTRL livers, cell area decreased and cell density increased across pregnancy (Fig. 2K-L). Conversely, NP LERKO livers exhibited reduced cell area and increased baseline cell density compared with CTRL livers (Fig. 2K-L). During pregnancy, cell area in LERKO livers remained relatively stable, whereas cell density increased at GD15 compared to the NP condition (Fig. 2K-L).

### Transcriptomics reveals dynamic waves of liver remodeling across gestation

Bulk RNA sequnecing of pregnant CTRL livers identified 236, 955, and 722 differentially expressed genes (DEGs) at GD7, GD15, and GD18, respectively, compared with NP CTRL (|FC| > 2, *p*-adj < 0.05; Fig. 3A and Fig. S1). Pairwise comparisons between GD15/GD7 and GD18/GD15 yielded 743 and 196 DEGs, respectively (Fig. 3A). Venn and UpSet analyses revealed both shared and stage-specific transcriptional signatures across gestation (Fig. 3B-C, Fig. S2–S3).

**Fig. 3.**
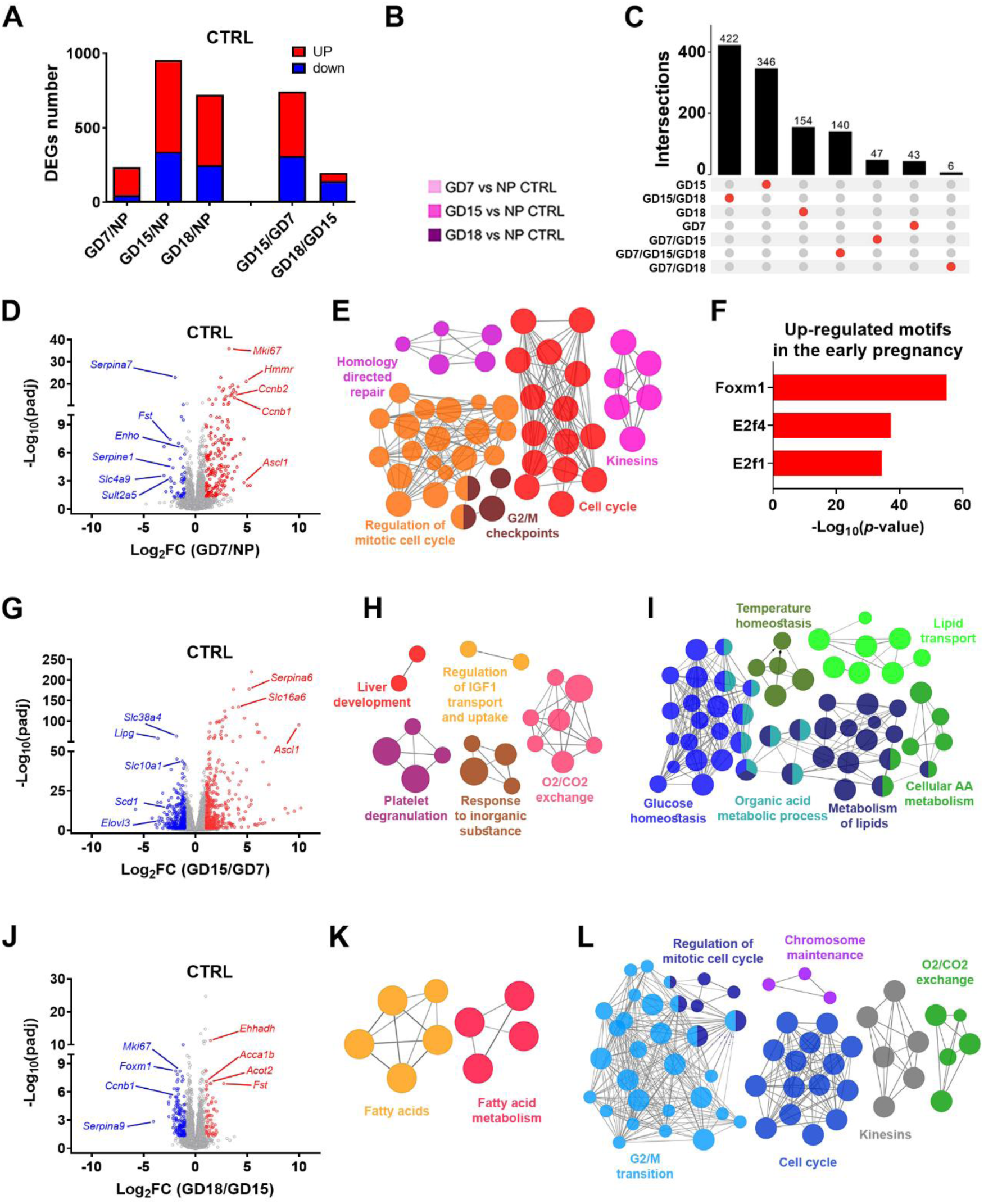
Pregnancy progression induces dynamic liver transcriptome remodeling. (**A**) Distribution of up- and down-regulated genes in CTRL liver at GD7, GD15, and GD18 relative to the NP condition, as determined by RNA-seq. (**B-C**) Venn diagram (**B**) and UpSet plot (**C**) showing shared and stage-specific liver DEGs at GD7, GD15 and GD18 compared with NP CTRL. (**D**) Volcano plot of liver DEGs in early pregnancy (GD7). (**E-F**) Gene Ontology (GO) analysis of functional networks (**E**) and enriched motifs (**F**) among genes upregulated in GD7 CTRL liver. (**G-I**) Volcano plot (**G**) and GO analysis of functional networks significantly up- (**H**) and down-regulated (**I**) in CTRL liver at GD15 compared with GD7. (**J-L**) Volcano plot (**J**) and GO analysis of functional networks significantly up- (**K**) and down-regulated (**L**) in CTRL liver at GD18 compared with GD15. RNA-seq analyses included n=3 independent CTRL animals at each reproductive stage.

At GD7, the hepatic transcriptome was enriched with genes associated with cell-cycle progression, mitotic activity, and DNA replication, including *Ccna2*, *Ccnb1*, *Ccnb2*, *Foxm1*, and *Mki67* (Fig. 3D– E, Fig. S2D). Motif enrichment analysis identified FOXM1 among the top predicted transcriptional regulators, consistent with its established role in hepatocyte proliferation and liver regeneration [33] (Fig. 3F). Additional upregulated genes included *Hmmr*, implicated in periportal hepatocyte expansion and glucose homeostasis during pregnancy [6], and *Ascl1*, a neurogenic factor essential for maternal liver adaptation [34] (Fig. 3D). Notably, 89 DEGs were commonly upregulated across GD7, GD15, and GD18, peaking at mid-pregnancy and declining thereafter (Fig. S2A, S2C), consistent with temporally regulated activation. Twenty-seven genes were uniquely induced at GD7, including those involved in lipid and fatty acid metabolism (i.e. *Elovl6*, *Acot3*), bile acid metabolism (i.e. *Cyp7a1*, *Cyp8b1*), steroid metabolism (i.e. *Srd5a1, Ugt2b37*), and glucose and lipid regulation (*Mup3* and *Zbtb16*) (Fig. S3A-C). Downregulated genes at GD7 were fewer and predominantly associated with immune regulation (*Cd79a, Pax5*), hepatokine signaling (*Enho, Fst, Serpine1*), and solute transport (*Slc4a9, Serpina7*), consistent with early immune and metabolic remodeling during pregnancy adaptation (Fig. 3D, Fig. S2H, S3D).

At GD15, both the number and magnitude of transcriptomic changes intensified (Fig. 3A, 3G). Key proliferation-associated genes and pregnancy-associated regulators, including *Ascl1*, *Hmmr*, and *Cyp3a44*, were markedly enhanced, consistent with the progression of the proliferative program initiated earlier in pregnancy (Fig. 3G-H, Fig. S1D, S2D, S4A). Genes involved in angiogenesis, extracellular matrix remodeling, solute transport, and hormone responsiveness were also induced at this stage (Fig. S4A–B). The peak in proliferation-associated gene expression at GD15 coincided with the downregulation of key metabolic genes, particularly those linked to AA catabolism, glucose metabolism, lipogenesis, lipid transport, and fatty acid oxidation (FAO) (Fig. 3I, Fig. S4C–F, S1H). Many of these genes are established targets of PPARα, which was downregulated at this stage (Fig. S4G).

By GD18, the hepatic transcriptome showed renewed expression of PPARα target genes involved in FAO, glucose metabolism, and hepatokine production, including *Cyp4a10*, *Cyp4a14*, *G6pc*, *Pdk4*, *Fgf21*, *Fst* (Fig. 3J-K, Fig. S1H). This late-gestation profile coincided with reduced expression of cell-cycle genes, including *Ccna2*, *Ccnb1*, *Ccnb2*, *Foxm1*, *Mki67* (Fig. 3J, 3L, Fig. S2D), consistent with reduced proliferation-associated transcriptional activity and restoration of selected metabolic programs (Fig. S1I).

Collectively, these findings indicate that the liver undergoes stage-specific transcriptional remodeling throughout pregnancy. This process is characterized by early induction of cell-cycle and proliferation-associated programs, a mid-gestational peak in proliferation-related gene expression coupled with suppression of selected metabolic pathways, and late reactivation of metabolic programs. These transcriptomic patterns were paralleled by dynamic changes in targeted liver metabolite profiles, with the largest shifts observed during early and mid-pregnancy (Fig. S5 and Supplementary Table S2).

### Hepatic ERα deficiency alters early proliferative and metabolic adaptation during pregnancy

To elucidate the impact of hepatic ERα deficiency on gestational liver adaptation, we first compared liver gene expression between CTRL and LERKO females under NP conditions. This analysis revealed that hepatic ERα deficiency already altered basal transcriptional programs, with reduced expression of differentiation-, metabolic-, and estrogen-responsive genes and increased expression of genes associated with proliferative, lipid-handling, and immune-related pathways (Fig. S6). Throughout gestation, the number of pregnancy-regulated DEGs was lower in LERKO compared to CTRL livers, except at GD7, when the most significant genotype-dependent transcriptional divergence was observed (Fig. 4A–B).

**Fig. 4.**
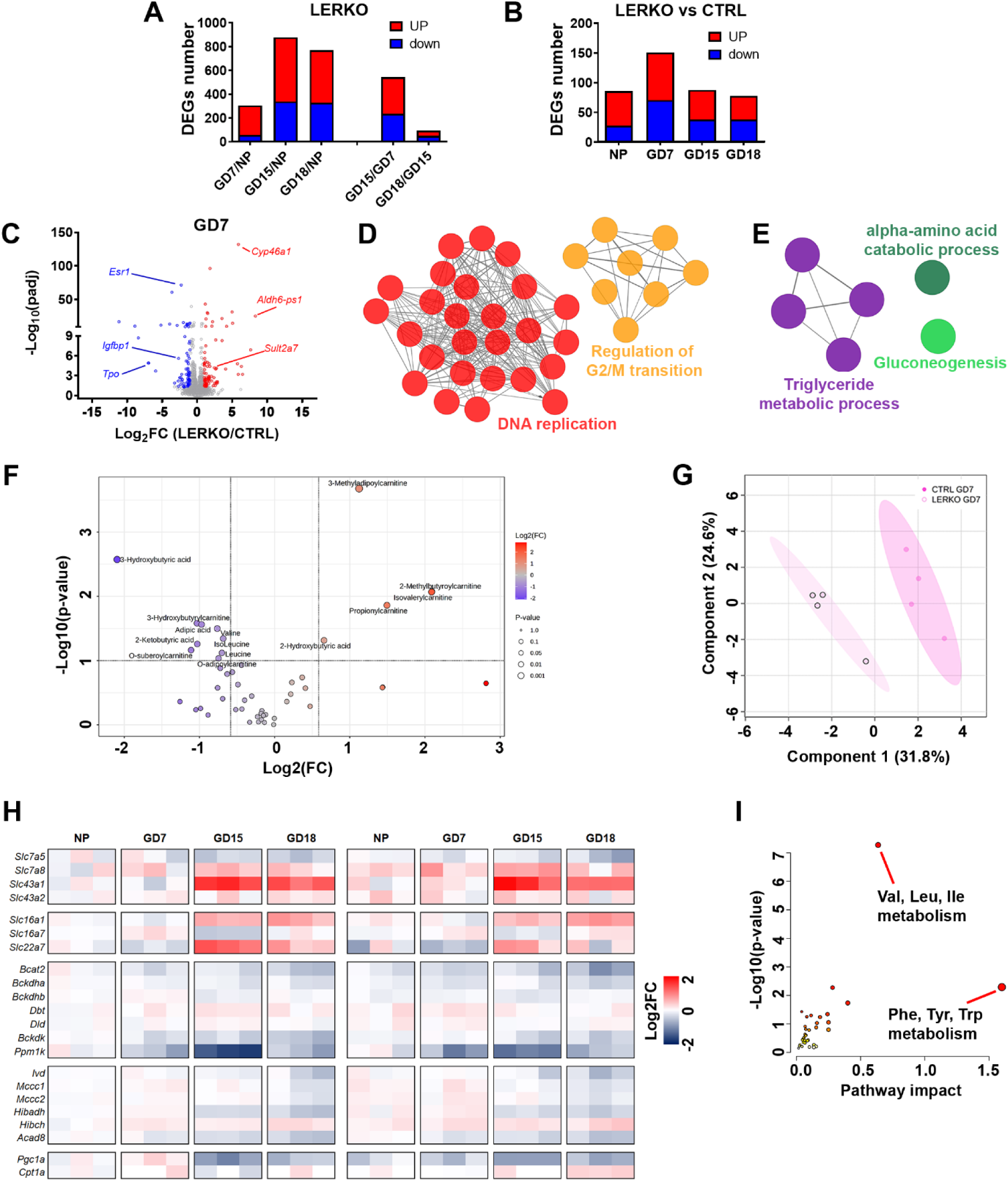
Hepatic ERα deficiency alters early pregnancy adaptation. (**A**) Distribution of up- and down-regulated genes in LERKO liver at GD7, GD15, and GD18 relative to the NP condition, as determined by RNA-seq. (**B**) DEGs in LERKO liver compared with CTRL liver across gestation. (**C-E**) Volcano plot (**C**) and GO analysis of significantly up- (**D**) and down-regulated (**E**) functional networks in LERKO liver compared with CTRL at GD7. (**F-G**) Volcano plot (**F**) and principal component analysis (PCA; **G**) of selected metabolites in CTRL (n=4) and LERKO (n=4) livers at GD7. (**H**) Heatmap showing hepatic expression of genes involved in BCAA/BCKA handling and catabolism, mitochondrial oxidative metabolism, and fatty acid oxidation in CTRL and LERKO females across gestation. (**I**) Integrated metabolomic and transcriptomic analysis showing pathways altered in LERKO compared with CTRL liver at GD7. RNA-seq analyses included n=3 independent animals per genotype and reproductive stage.

At GD7, LERKO livers displayed 143 uniquely regulated genes compared to CTRL livers (Fig. S7A-J), including the upregulation of cell-cycle and DNA replication/repair genes such as *Mcm2-7, Pcna,* and *Pole* (Fig. 4D, Fig. S8A). At the same time, genes involved in glycolysis (*Pklr*) and FA synthesis (*Fasn*) were upregulated, whereas genes related to AA catabolism (*Got1, Sds, Tat*), gluconeogenesis (*G6pc, Pck1*), and PPARα signaling (*Cyp8b1, Plin4*) were downregulated (Fig. 4D-E, Fig. S8B–E). Additionally, key regulatory transcripts involved in hormonal and energy sensing (*Enho* and *Igfbp1*) were reduced in LERKO livers (Fig. S8F). Collectively, these changes indicate altered regulation of proliferative, anabolic, and metabolic transcriptional programs in LERKO livers during early pregnancy.

Targeted metabolomic profiling was consistent with these transcriptional changes. At GD7, LERKO livers showed reduced levels of β-hydroxybutyrate (β-BHB) and branched-chain amino acids (BCAA), along with accumulation of acylcarnitines, consistent with altered handling of FA β-oxidation intermediates and AA catabolism (Fig. 4F–G). In line with this profile, transcripts encoding components of the BCAA/BCKA oxidative pathway, the BCKA–;related transporter *Slc22a7*, and mitochondrial metabolic regulators including *Cpt1a* and *Ppargc1a* were reduced in LERKO livers at GD7 (Fig. 4H). Integrated multi-omics analysis identified AA metabolism as the most affected pathway during early pregnancy adaptation in LERKO livers (Fig. 4I).

In summary, these data underscore support the role of hepatic ERα in the proper organization of early proliferative and metabolic adaptations during pregnancy. Its absence was associated with premature induction of proliferative and anabolic transcriptional programs and altered AA- and FA-related metabolic signatures, consistent with an early mismatch between liver growth programs and metabolic adaptation.

### Loss of hepatic ERα alters mid-gestational proliferative, metabolic, and insulin-related responses

Next, we focused on GD15, a stage at which LERKO females exhibited a higher liver-to-body weight *ratio* than CTRL mice (Fig. 2). In CTRL livers, GD15 was associated with a significant induction of mitotic genes, whereas LERKO livers showed a weaker induction of these markers, indicating a reduced proliferative response at mid-gestation (Fig. 5A-D). Genes involved in DNA replication initiation and repair, such as *Mcm5* and *Mcm6*, which were transiently upregulated in LERKO livers at GD7, were no longer elevated at GD15 (Fig. S8A, S9G). This pattern is consistent with premature activation of cell-cycle-associated programs in LERKO livers, followed by a reduced proliferative response at the stage when liver growth typically peaks. Ki67 immunostaining supported this interpretation, showing a lower percentage of Ki67-positive nuclei in LERKO livers, with the largest genotype-dependent difference observed at GD15 (Fig. 5E–F).

**Fig. 5.**
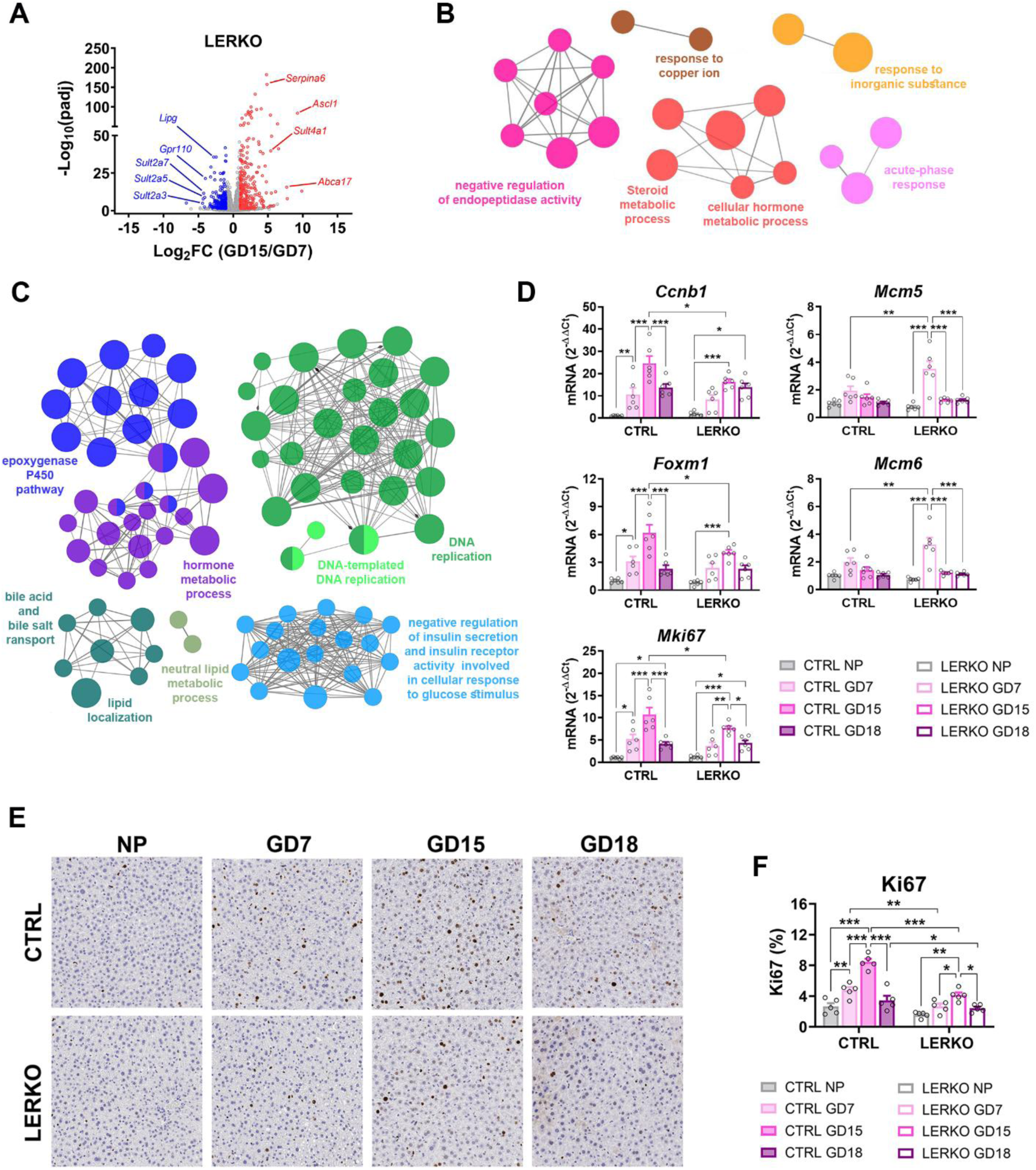
Hepatic ERα deficiency alters mid-gestational proliferative responses. (A-C) Volcano plot (**A**) and GO analysis of biological processes associated with genes significantly up- (**B**) and down-regulated (**C**) in LERKO liver at GD15 compared with GD7 by RNA-seq (n=3 per group). (**D**) mRNA levels of *Ccnb1*, *Foxm1*, *Mki67*, *Mcm5* and *Mcm6* measured by qPCR in CTRL and LERKO livers across reproductive stages (n=6 per group). (**E-F**) Representative images (**E**) and quantification (**F**) of Ki67-positive nuclei (%) in CTRL and LERKO livers across reproductive stages (n=5 per group). Data are mean ± SEM; *p<0.05, **p<0.01 and ***p<0.001 by two-way ANOVA followed by Bonferroni’s *post hoc* test.

The transition from GD7 to GD15 in LERKO livers was also characterized by decreased expression of genes involved in metabolic and endocrine-related pathways, including the epoxygenase P450 pathway, hormone metabolism, bile acid transport, and insulin signaling, such as *Insrr*, *Irs1,* and *Mup11/12/16/2* (Fig. 5C). These findings are consistent with altered regulation of metabolic and endocrine-related transcriptional programs during mid-gestation in LERKO livers.

To assess whether these hepatic alterations were accompanied by changes in systemic glucose handling, we performed glucose (GTT) and insulin (ITT) tolerance tests. Compared with non-pregnant CTRL mice, GD15 CTRL mice showed an attenuated glucose excursion, consistent with normal gestational adaptation, whereas LERKO mice displayed a pronounced glucose peak followed by delayed clearance (Fig. 6A–B). ITT responses were comparable between genotypes (Fig. 6C-D), suggesting that altered glucose handling in LERKO mice occurred without overt systemic insulin resistance. Consistent with this interpretation, circulating glucose and insulin levels were similar between groups (Fig. S10). Hepatic glycogen content, however, differed significantly at GD7, consistent with altered hepatic glycogen handling early in pregnancy (Fig. S10).

**Fig. 6.**
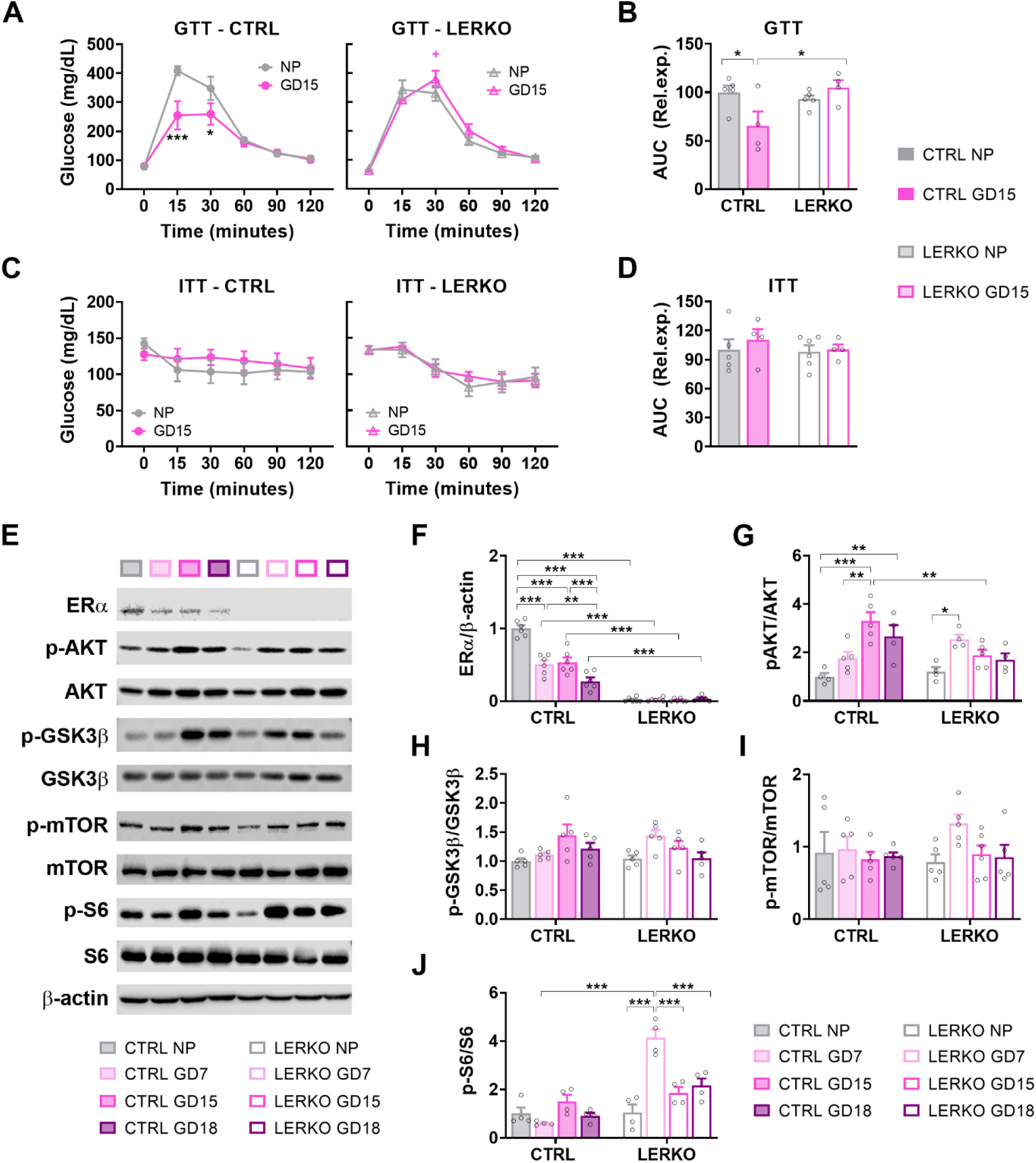
LERKO females exhibit altered glucose handling and AKT–mTORC1-related signaling at mid-gestation. (**A-B**) Blood glucose profile (**A**) and area under the curve (AUC; **B**) during the glucose tolerance test (GTT) in NP and GD15 CTRL and LERKO mice (CTRL NP, n=5; CTRL GD15, n=4; LERKO NP, n=5; LERKO GD15, n=4). (**C-D**) Blood glucose profile (**C**) and AUC (**D**) during the insulin tolerance test (ITT) in NP and GD15 CTRL and LERKO (CTRL NP, n=5; CTRL GD15, n=4; LERKO NP, n=5; LERKO GD15, n=4). (**E**-**J**) Representative western blots (**E**) and semi-quantitative analyses of ERα (**F**; n=6 per group), p-AKT (Ser473)/AKT (**G**; CTRL NP/GD7/GD15/GD18, n=4/5/5/4; LERKO NP/GD7/GD15/GD18, n=4/4/5/4), p-GSK3β (Ser9)/GSK3β (**H**; n=5 per group), p-mTOR (Ser2448)/mTOR (**I**; n=5 per group), and p-S6 (Ser235/236)/S6 (**J**; n=4 per group) measured in liver extracts from NP and pregnant CTRL and LERKO mice. Data are mean ± SEM; *p<0.05, **p<0.01 and ***p<0.001 by two-way ANOVA followed by Bonferroni’s *post hoc* test.

At the molecular level, targeted analysis of basal hepatic signaling indicated altered temporal regulation of AKT-related signaling in LERKO mice. In CTRL livers, the pAKT/AKT *ratio* increased during pregnancy, peaking at GD15. In contrast, LERKO livers showed an earlier increase at GD7 but did not reach CTRL levels at GD15, consistent with altered timing of AKT signaling (Fig. 6E– G). The pGSK3β/GSK3β *ratio* followed broadly similar patterns across gestation in the two genotypes (Fig. 6E, 6H).

Given the central role of mTORC1 in proliferation and metabolism [35,36], and its crosstalk with ERα and AKT [37,38], we next examined downstream mTORC1-related signaling. The p-mTOR/mTOR *ratio* remained relatively stable throughout gestation, whereas an increased pS6/S6 *ratio* in LERKO livers at GD7 suggested an early alteration in selected downstream mTORC1-related signaling markers (Fig. 6E, 6I-J).

To further assess insulin responsiveness, mice were subjected to an acute insulin challenge (Fig. 7). In CTRL livers, insulin induced phosphorylation of AKT, GSK3β, and S6 in NP and GD7 animals, whereas S6K1 phosphorylation was evident in NP but not at GD7. AKT responsiveness was reduced at GD15, consistent with physiological modulation during pregnancy. In this exploratory experiment, LERKO livers showed a pattern of maintained insulin-induced S6K1 and S6 phosphorylation at GD15 without a corresponding genotype-dependent difference in proximal AKT activation (Fig. 7), suggesting a potential alteration in the coupling between proximal AKT and downstream signaling. Collectively, these findings suggest that loss of hepatic ERα is associated with altered timing of proliferative responses, altered gestational glucose adaptation, and changes in basal AKT–mTORC1-related signaling, along with preliminary evidence of altered insulin-responsive signaling at GD15. Specifically, LERKO livers showed an early alteration in selected downstream anabolic signaling markers at GD7, followed by altered proliferative and metabolic adaptation at GD15.

**Fig. 7.**
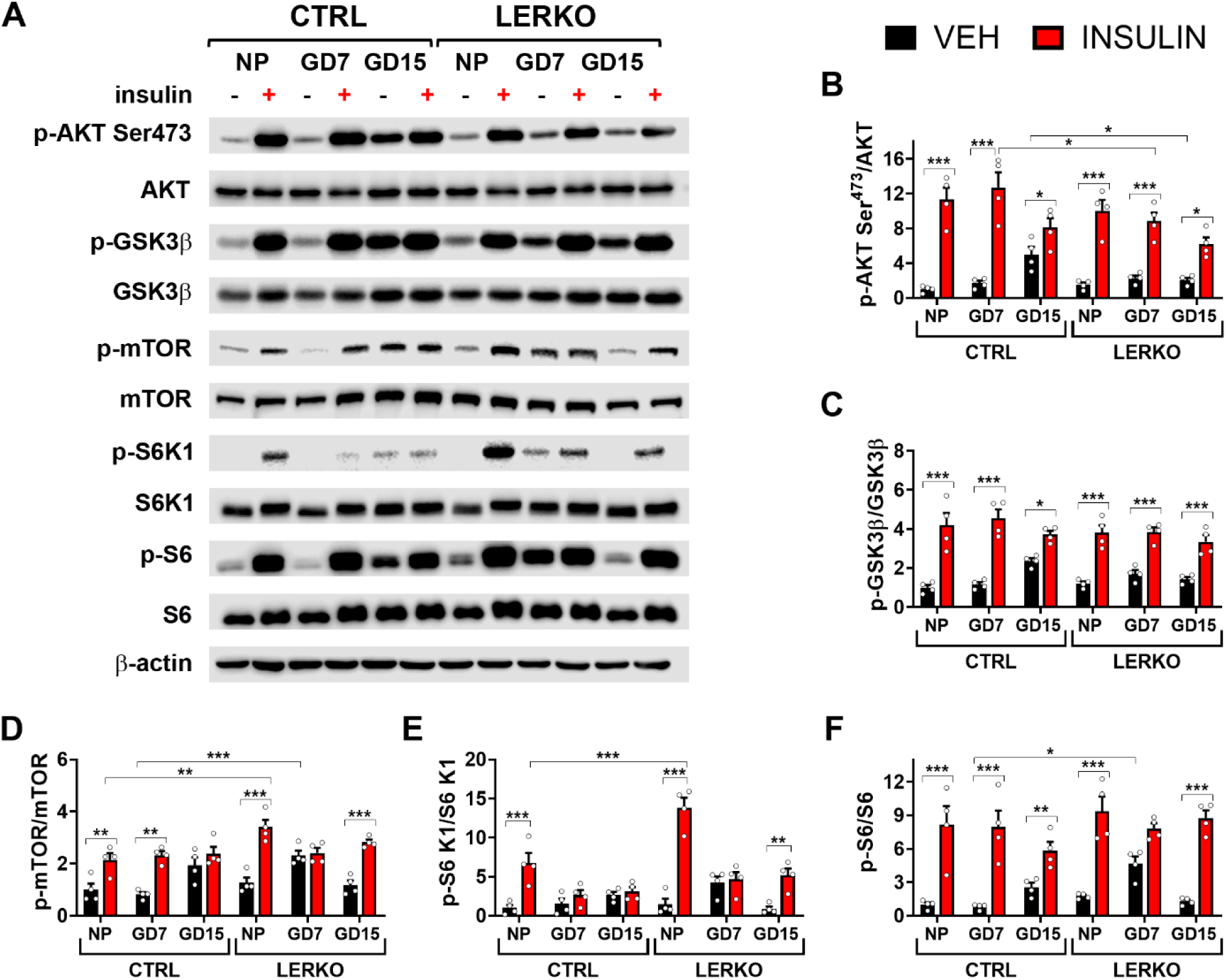
Exploratory analysis of insulin-induced AKT–mTORC1-related signaling during pregnancy in hepatic ERα-deficient mice. (**A-F**) Representative western blots (**A**) and semi-quantitative analyses of total and phosphorylated AKT (Ser473) (**B**), GSK3β (Ser9) (**C**), mTOR (Ser2448) (**D**), S6K1 (Thr389) (**E**), and S6 (Ser235/236) (**F**) in liver from NP, GD7, and GD15 CTRL and LERKO female mice following vehicle or insulin treatment. Data are mean ± SEM (n=4 biological replicates per genotype × reproductive stage × treatment subgroup); *p<0.05, **p<0.01 and ***p<0.001 by genotype × treatment two-way ANOVA performed separately within each reproductive stage, followed by Bonferroni-adjusted comparisons.

### Pregnancy is associated with altered light/dark-phase energy-balance patterns in LERKO females

To assess systemic energy-balance patterns associated with hepatic ERα deficiency at mid-pregnancy, we performed indirect calorimetry in NP and pregnant CTRL and LERKO females. Pregnancy resulted in increased oxygen consumption (O₂) and carbon dioxide production (CO₂), reflecting heightened metabolic demands during gestation. In comparison to CTRL mice, CO_2_ production was significantly elevated in NP LERKO mice during the dark phase and in pregnant LERKO mice during the light phase (Fig. 8A–B).

**Fig. 8.**
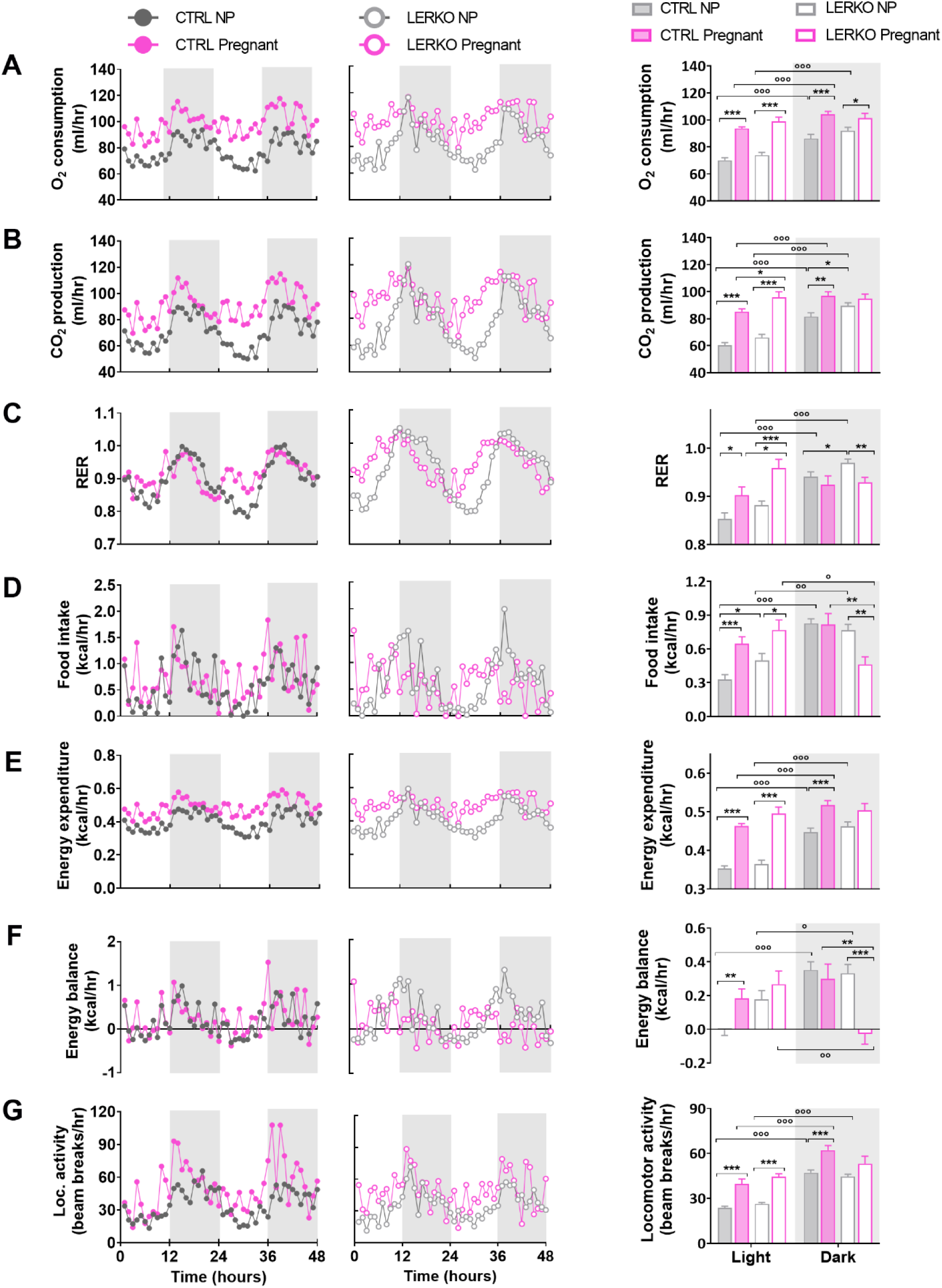
Hepatic ERα deficiency alters light/dark-phase energy-balance patterns at mid-gestation. (**A-G**) Hourly profiles and phase-averaged values for O_2_ consumption (**A**), CO_2_ production (**B**), respiratory exchange *ratio* (RER; **C**), food intake (**D**), energy expenditure (**E**), energy balance (**F**), and locomotor activity (**G**) measured in CTRL and LERKO by indirect calorimetry. Light and dark phases are distinguished by white and gray backgrounds, respectively. Data are mean ± SEM (CTRL NP, n=7; CTRL pregnant, n=5; LERKO NP, n=7; LERKO pregnant, n=6). Statistical analyses were performed on phase-averaged data using a mixed-design repeated-measures ANOVA, with experimental group (CTRL NP, CTRL GD15, LERKO NP, and LERKO GD15) as the between-subject factor and light/dark phase as the within-subject factor, followed by Bonferroni-adjusted prespecified comparisons. *p<0.05, **p<0.01 and ***p<0.001 for genotype comparisons within the indicated reproductive condition and phase; °p<0.05, °°p<0.01 and °°°p<0.001 for dark-*versus* light-phase comparisons within the indicated experimental group.

LERKO mice also exhibited higher respiratory exchange *ratio* (RER) compared with CTRL mice. Specifically, RER was increased in NP LERKO during the dark phase, whereas pregnant LERKO mice showed higher RER during the light phase (Fig. 8C). These findings are compatible with altered substrate utilization, including a relative shift toward carbohydrate utilization, in line with the hepatic transcriptional evidence of altered FAO and PPARα-associated pathways.

Feeding behavior also differed between genotypes: NP LERKO showed increased food intake during the light phase, whereas pregnant LERKO exhibited reduced food intake during the dark phase compared with CTRL mice (Fig. 8D). Energy expenditure was higher during the dark phase in all groups except pregnant LERKO mice (Fig. 8E). Accordingly, pregnant LERKO mice displayed a positive energy balance during the light phase and a negative energy balance during the dark phase (Fig. 8F). Locomotor activity was greater during the dark than the light phase in all groups except pregnant LERKO mice (Fig. 8G).

No major genotype-specific differences in white adipose tissue mass or gene expression were observed (Fig. S11). As hepatic clock genes were measured only at a single experimental time point (Fig. S12), these findings should be interpreted as indicative of altered light/dark-phase metabolic organization rather than direct evidence of disruption of the core circadian clock machinery.

## Discussion

Pregnancy requires dynamic coordination of maternal liver growth, substrate metabolism, and endocrine signaling. Here, we show that hepatic ERα contributes to the stage-specific organization of these processes during gestation. Previous work from our group and others has identified hepatic ERα as a regulator of lipid oxidation, amino acid metabolism, hepatokine expression, and PPARα-associated pathways in the non-pregnant female liver [10,11,13–15]. The current study advances this understanding by revealing a temporal dimension to hepatic ERα function: it is essential not only for regulating individual metabolic pathways but also for aligning proliferative, metabolic, and insulin-responsive programs throughout gestation.

In control mice, liver adaptation followed a sequential pattern, with early induction of cell-cycle-related programs, a mid-gestational proliferative peak coupled with the suppression of selected metabolic pathways, and late reactivation of metabolic programs. In LERKO mice, this temporal organization was disrupted. Early pregnancy was associated with premature induction of proliferative and anabolic transcriptional programs and altered AA- and FA-related metabolic signatures, whereas mid-gestation was characterized by reduced proliferation-associated gene expression, lower Ki67-positive nuclei, altered insulin-related transcriptional programs, and altered gestational glucose adaptation. Thus, hepatic ERα deficiency appears to uncouple the timing of liver growth programs from metabolic adaptation. Because the metabolomic analyses were steady-state and flux-based assays were not performed, these findings should be interpreted as indicative of altered substrate handling rather than direct evidence of impaired mitochondrial oxidation.

The altered AKT-mTORC1-related signaling observed in LERKO livers provides a plausible molecular correlate of this temporal mismatch. Basal analyses indicated an early alteration in selected downstream mTORC1-related signaling markers in LERKO livers at GD7 (Fig. 6). In the acute insulin-stimulation experiment, LERKO livers showed sustained insulin-induced S6K1/S6 phosphorylation at GD15 despite comparatively modest proximal AKT activation (Fig. 7). Although preliminary, this pattern suggests altered coupling between proximal insulin-AKT signaling and downstream anabolic responses, a pathway axis that may influence both hepatocyte proliferation and metabolic remodeling.

Systemic phenotyping further supports the view that hepatic ERα deficiency is associated with altered gestational metabolic adaptation. GTT and ITT analyses were performed at GD15 because this stage coincided with impaired hepatocyte proliferation, altered AKT–mTORC1-related signaling, and the emergence of impaired glucose handling in LERKO dams. The comparable ITT responses and preserved circulating insulin levels argue against generalized systemic insulin resistance but instead suggest altered gestational glucose handling associated with hepatic ERα deficiency. By contrast, GD7 represented an early phase characterized by hepatic molecular and metabolic alterations, including altered glycogen handling and AA- and FA-related signatures. Whether these early hepatic changes are already sufficient to impair systemic glucose or insulin tolerance remains unresolved.

Indirect calorimetry revealed altered light/dark-phase patterns of substrate utilization, feeding behavior, energy expenditure, and locomotor activity during the light/dark phases in pregnant LERKO mice. These observations suggest a modification in diurnal metabolic organization; however, they do not confirm a disruption of the core circadian clock, as hepatic clock genes were evaluated at only a single time point. Furthermore, systemic metabolic phenotypes observed during pregnancy may represent an integration of maternal, fetal, and placental physiology rather than autonomous effects to the liver.

These findings have implications for pregnancy-associated metabolic and liver disorders, including GDM, MASLD, and ICP [22–27]. Although these conditions are multifactorial and direct extrapolation from mice to humans remains speculative, it remains unknown whether the hepatic transcriptomic and metabolomic alterations observed in LERKO mice are mirrored in human pregnancy-associated metabolic or liver disease.

These findings may pregnancy-associated metabolic and liver disorders, such as gestational diabetes mellitus (GDM), metabolic-associated steatotic liver disease (MASLD), and intrahepatic cholestasis of pregnancy (ICP) [22–27]. Although these conditions are multifactorial and direct extrapolation from mice to humans remains speculative, it is yet to be determined whether the hepatic transcriptomic and metabolomic changes observed in LERKO mice are reflected in human pregnancy-associated metabolic or liver diseases.

The present study identifies hepatic ERα signaling as a potential determinant of maternal liver adaptability during gestation. In this framework, impaired hepatic ERα activity may not simply alter individual metabolic pathways but may compromise the temporal coordination between liver growth, substrate utilization, and insulin-responsive signaling under pregnancy-associated metabolic stress. Future human genetic, transcriptomic, metabolomic, and clinical studies will be required to ascertain whether similar ERα-dependent adaptive programs are conserved in human pregnancy and altered in pregnancy-associated liver disease.

## Conclusions

In conclusion, our findings support a requirement for hepatic ERα in the appropriate stage-specific coupling of metabolic, proliferative, and insulin-responsive programs during pregnancy. These results extend the established role of hepatic ERα in female liver metabolism to physiological adaptation during gestation and provide a framework for future mechanistic and translational studies.

## Limitations of the study

While this study supports a requirement for hepatic ERα in stage-specific metabolic–proliferative coupling during pregnancy, several limitations should be acknowledged. This study does not distinguish direct ERα transcriptional targets from indirect downstream effects, as ERα chromatin occupancy was not assessed in the pregnant liver. Thus, transcriptomic changes may reflect primary ERα-regulated events or secondary outcomes of altered metabolic states, signaling activities, intercellular crosstalk, or chronic adaptation to ERα deficiency. The constitutive Albumin-Cre LERKO model cannot separate acute gestational functions of ERα from developmental or long-term compensatory effects, given that LERKO females enter pregnancy with a chronically ERα-deficient and potentially pre-adapted liver. Future studies using temporally controlled hepatocyte-specific ERα perturbation approaches (e.g., AAV8-TBG-Cre or inducible Cre systems) will be required to address this distinction.

Rescue experiments were not performed. Although the data support a requirement for hepatic ERα in gestational liver adaptation, they do not confirm whether ERα re-expression or targeted modulation of downstream pathways such as AKT-mTORC1 would restore normal proliferative and metabolic remodeling.

Whole-liver bulk RNA-seq does not resolve cell-type-specific contributions. Although Albumin-Cre primarily targets hepatocytes, non-parenchymal cells may contribute to pregnancy-associated hepatic remodeling through paracrine, immune, vascular, or stromal mechanisms. Single-cell or spatial approaches will be needed to distinguish hepatocyte-autonomous effects from intercellular contributions.

Metabolomic analyses were based on targeted steady-state measurements and did not include comprehensive lipidomics, isotope tracing, or flux-based assays. Therefore, conclusions regarding lipid remodeling, FAO, AA catabolism, gluconeogenesis, and substrate fluxes should be interpreted as inferential rather than direct measures of pathway activity.

Systemic and translational interpretations remain limited. GTT and ITT were performed only in NP and GD15 pregnant mice; future studies with larger pregnancy cohorts will be required to define the temporal onset of whole-body glucose-handling alterations in LERKO dams.

Given that LERKO dams showed increased litter-size variability, reduced fetal weight, and a trend toward fetal resorption, mid- and late-gestational metabolic phenotypes may also reflect altered fetal or placental demand or resorption-associated responses.

Indirect calorimetry in pregnant mice may be influenced by single housing and metabolic cage stress; corticosterone was not measured, and altered light/dark-phase metabolic patterns cannot be interpreted as core circadian clock disruption because clock genes were assessed at only one time point.

Finally, this study was conducted exclusively in mice, and future human studies will be required to determine whether similar ERα-dependent adaptive programs are conserved in human pregnancy and altered in pregnancy-associated metabolic disorders.

## Resource availability

### Data availability

Raw RNA-seq data have been deposited in BioProject. Processed metabolomic data and uncropped western blot images are provided as Supplementary Data.

## Acknowledgements

We are grateful to Dr. Valeria Benedusi for English proofreading and critical discussion, and to DISFARM for administrative support. This study was supported in part by European Union’s Horizon Europe Research and Innovation Programme (Ref. 101080329) to A.G.; the European Community (ERC-Advanced Grant 322977) to A.M; EMBO Short-term Fellowship (Ref. ASTF 184-2016), DAAD Award (Ref. 91799341) and Pfizer Global NASH ASPIRE Competitive Grant Program (Grant 77230651) to S.D.T.

## Author Contributions

Conceptualization: S.D.T. Data curation: S.D.T. Formal analysis: S.D.T. Funding acquisition: S.D.T., A.M., and A.G. Investigation: S.D.T., A.D., C.M., G.T., F.C., and P.I. Project administration: C.M. and S.D.T. Resources: S.D.T., C.M., A.G., and C.O. Supervision: S.D.T. Visualization: S.D.T. Writing - original draft: S.D.T. Writing - review and editing: all authors.

## Declaration of interests

A.G. reports consulting or advisory roles for Boehringer Ingelheim, Eli Lilly and Company, Metadeq Diagnostics, Merck Sharp & Dohme, Novo Nordisk, and Pfizer, and speaker honoraria or other fees from Eli Lilly and Company, Merck Sharp & Dohme, Novo Nordisk, Madrigal, Echosens, Mercodia, and Pfizer. The other authors declare no competing interests.

## Declaration of AI-assisted technologies in the manuscript preparation process

During the preparation of this work, the authors used ChatGPT and Claude to improve the English language and readability of the manuscript. The authors reviewed and edited the output as needed and take full responsibility for the content of the published article.

