## Supplementary information for "Hepatic estrogen receptor α is required for stage-specific coupling of liver metabolism and proliferation during pregnancy"

**Abbreviations of genes cited in the manuscript and figures**

*Abcb4,* ATP binding cassette subfamily B member 4

*Abcc3*, ATP binding cassette subfamily C member 3

*Acaa1b,* acetyl-Coenzyme A acyltransferase 1B

*Acot1,* acyl-CoA thioesterase 1

*Acot2,* acyl-CoA thioesterase 2

*Acot3*, acyl-CoA thioesterase 3

*Adipoq*, adiponectin

*Agpat2,* 1-acylglycerol-3-phosphate O-acyltransferase 2

*Arntl,* aryl hydrocarbon receptor nuclear translocator-like

*Ascl1*, achaete-scute family bHLH transcription factor 1

*Asl,* argininosuccinate lyase

*Ass1,* argininosuccinate synthase 1

*Atgl* (or *Pnpla2*), patatin-like phospholipase domain containing 2

*Bcat2,* branched chain amino acid transaminase 2

*Bckdha,* branched chain keto acid dehydrogenase E1 subunit alpha

*Ccna2,* cyclin A2

*Ccnb1,* cyclin B1

*Ccnb2,* cyclin B2

*Cd79a,* CD79a molecule

*Clock,* clock circadian regulator

*Cpt1α,* carnitine palmitoyltransferase 1a

*Cry1*, cryptochrome circadian regulator 1

*Cry2*, cryptochrome circadian regulator 2

*Cth,* cystathionine gamma-lyase

*Cyp3a44,* cytochrome P450, family 3, subfamily a, polypeptide 44

*Cyp4a10,* cytochrome P450, family 4, subfamily a, polypeptide 10

*Cyp4a14,* cytochrome P450, family 4, subfamily a, polypeptide 14

*Cyp4f15,* cytochrome P450, family 4, subfamily f, polypeptide 15

*Cyp7a1,* cytochrome P450 family 7 subfamily A member 1

*Cyp8b1,* cytochrome P450 family 8 subfamily B member 1

*Cyp46a1*, cytochrome P450 family 46 subfamily A member 1

*Ehhadh,* enoyl-CoA hydratase and 3-hydroxyacyl CoA dehydrogenase

*Elovl6,* ELOVL fatty acid elongase 6

*Enho,* energy homeostasis associated

*Fasn,* fatty acid synthase

*Fgf21,* fibroblast growth factor 21

*Foxm1,* forkhead box M1

*Fst,* follistatin

*G6pc,* glucose-6-phosphatase catalytic subunit

*Ghr,* growth hormone receptor

*Gls2,* glutaminase 2

*Got1,* glutamic-oxaloacetic transaminase 1

*Hmmr,* hyaluronan mediated motility receptor

*Hsd11b1,* hydroxysteroid 11-beta dehydrogenase 1

*Hsl* (also *Lipe*), lipase, hormone sensitive

*Igfbp1,* insulin like growth factor binding protein 1

*Inhbb,* inhibin subunit beta B

*Insrr,* insulin receptor related receptor

*Irs1,* insulin receptor substrate 1

*Ki67,* marker of proliferation Ki-67

*Lepr,* leptin receptor

*Lipc,* lipase C, hepatic type

*Lipg,* lipase G, endothelial type

*Lep,* leptin

*Mcm2,* minichromosome maintenance complex component 2

*Mcm3,* minichromosome maintenance complex component 3

*Mcm4,* minichromosome maintenance complex component 4

*Mcm5,* minichromosome maintenance complex component 5
*Mcm6,* minichromosome maintenance complex component 6
*Mcm7,* minichromosome maintenance complex component 7
*Mup2,* major urinary protein 2
*Mup3,* major urinary protein 3
*Mup11*, major urinary protein 11
*Mup12*, major urinary protein 12
*Mup16*, major urinary protein 16
*Mup21,* major urinary protein 21
*Nr1d1*, nuclear receptor subfamily 1 group d member 1
*Nr1d2*, nuclear receptor subfamily 1 group d member 2
*Npc,* NPC intracellular cholesterol transporter 1
*Pax5,* paired box 5
*Pck1,* phosphoenolpyruvate carboxykinase 1
*Pcna,* proliferating cell nuclear antigen
*Pdk4,* pyruvate dehydrogenase kinase 4
*Per1*, period circadian regulator 1
*Per2*, period circadian regulator 2
*Per3*, period circadian regulator 3
*Prlr*, prolactin receptor
*Pgc1α*, PPARG coactivator 1 alpha
*Pklr,* pyruvate kinase L/R
*Plin4,* perilipin 4
*Pnpla5,* patatin like domain 5, triacylglycerol lipase
*Pole,* DNA polymerase epsilon, catalytic subunit
*Pparα,* peroxisome proliferator activated receptor alpha
*Pparγ,* peroxisome proliferator activated receptor gamma
*Rorα*, RAR related orphan receptor a
*Rplp0*, ribosomal protein lateral stalk subunit p0
*Scd1,* stearoyl-Coenzyme A desaturase 1
*Sds,* serine dehydratase
*Serpina6,* serpin family A member 6
*Serpina7,* serpin family A member 7
*Serpine1,* serpin family E member 1

*Sult4a1,* sulfotransferase family 4A member 1

*Slc4a9,* solute carrier family 4 member 9

*Slc10a1,* solute carrier family 10 member 1

*Slc16a6,* solute carrier family 16 member 6

*Slc22a5,* solute carrier family 22 member 5

*Slc37a1*, solute carrier family 37 member 1

*Slc41a2*, solute carrier family 41 member 2

*Slco1a1,* solute carrier organic anion transporter family, member 1a1

*Slco1a4,* solute carrier organic anion transporter family, member 1a4

*Slco1b2,* solute carrier organic anion transporter family, member 1b2

*Srd5a1,* steroid 5 alpha-reductase 1

*Tat,* tyrosine aminotransferase

*Timeless,* timeless circadian regulator

*Ugt2b37,* UDP glucuronosyltransferase 2 family, polypeptide B37

*Vldlr,* very-low density lipoprotein receptor

*Zbtb16*, zinc finger and BTB domain containing 16

### Supplementary Materials and Methods

#### Animals and experimental design

Syngeneic ERα floxed (CTRL) and LERKO female mice were both C57BL/6J strain [28]. Four- to five-month-old mice were fed *ad libitum* with a standard diet (Ssniff V1534-703), provided with filtered water, and maintained at 22–25°C, 50±10% relative humidity, under an automatic 12-h light/dark cycle. Females were mated in the *proestrus/estrus* phase, with gestational day (GD) 0 defined as the day of mating. Pregnant females were euthanized at GD7, GD15, and GD18, whereas non-pregnant (NP) females were collected during *proestrus* to provide a hormonally standardized baseline comparison. Mice were euthanized after 6-h fasting in the early afternoon to minimize feeding- and time-dependent metabolic variability [28]. For acute insulin stimulation, 6h-fasted mice received intraperitoneal human insulin (0.75 U/kg) or vehicle, and livers were collected 12 min later for signaling analyses. All protocols were approved by the Istituto Superiore di Sanità ethics committee (1272/2015-PR and 149/2022-PR) and followed ARRIVE guidelines and European regulations.

#### Sex hormone measurements

17β-estradiol, estrone, progesterone, androstenedione, testosterone, and DHT were quantified by validated gas chromatography-tandem mass spectrometry, with quantification limits of 0.5, 0.5, 74, 12, 8, and 2.5 pg/mL, respectively, as previously described [29].

#### Biochemical analysis

Triglycerides (TG) were measured using commercial kits according to the manufacturer's protocols (Abcam, ab65336). Plasma levels of insulin were measured with Ultra Sensitive Mouse Insulin ELISA Kit (Crystal Chem, #90080) according to the manufacturers' protocol. Liver content of glycogen was measured with Glycogen Assay Kit (Abcam, ab65620) according to the manufacturer's protocol.

#### RNA extraction and qPCR

Total liver RNA was isolated, reverse transcribed, and analyzed by qPCR as previously described [30]. Primer sequences are reported in Table S1. Gene expression was normalized to *Rplp0* and calculated using the $2^{-\Delta\Delta Ct}$ method [31].

### Library preparation and RNA-sequencing

RNA quality was assessed using the Agilent 4200 TapeStation System (Agilent) and samples with adequate RNA Integrity Number values were processed. RNA concentration was estimated spectrophotometrically using Eppendorf BioSpectrometer Fluorescence instrument. Sequencing libraries were prepared using the Illumina® Stranded mRNA Prep, Ligation protocol (Illumina) with an input of 500 ng of total RNA. Final libraries were validated and quantified with the HSD1000 ScreenTape on the 4200 TapeStation System. Pooled libraries were sequenced on the Illumina NovaSeq platform, producing 2 × 150bp paired-end reads.

### Transcriptomics data analysis

Raw sequencing reads were processed for quality control using FASTQC (v0.11.5) (http://www.bioinformatics.babraham.ac.uk/projects/fastqc/). Adapter sequences were removed using Cutadapt. Paired-end reads were mapped to the mouse reference genome (GRCm38 primary assembly Gencode M20 and the associated GTF annotation file) using the STAR aligner (v2.5.2a). Raw read counts were generated using the quantMode TranscriptomeSAM option. Aligned data were manipulated using Samtools (v1.1). In particular, we evaluated coverage across the gene body, transcriptome profile efficiency (percentage of reads mapping to exons), samples correlation matrices, and the number of detected genes in order to identify possible contaminations, mapping failures, and potential outliers. Gene expression was quantified using the quantmode GeneCounts option in STAR. The counts produced are equivalent to those generated by htseq-count with default parameters. Sample counts were merged into a single gene counts-matrix, which was used as input for differential expression analysis. Statistical analysis was performed using the DESeq2 package (v1.30.0), testing group versus group accordingly to the experimental design (Wald Test). Unless otherwise stated, a threshold of 0.05 was applied to False Discovery Rate (FDR)-adjusted p-values to identify differentially expressed genes (DEGs) for downstream analysis. Exploratory data analysis (clustering and principal component analysis, PCA) was performed using built-in functions in the DESeq2 package. Gene ontology (GO) and cluster analysis were performed using the Cytoscape *plug-in* ClueGO [39], Genesis [40], Enrichr [41], and ShinyGO 0.82 (https://bioinformatics.sdstate.edu/go/). Venn diagrams were generated using Bioinformatics & Evolutionary Genomics software (http://bioinformatics.psb.ugent.be/webtools/Venn/) and Venn Diagram Maker Online (https://www.meta-chart.com/venn). Upset plot was generated using Hiplot software (https://hiplot.cn/basic/upset-plot).

**Metabolomic analysis**

A targeted metabolomic analysis was performed on approximately 25 mg of liver tissue. The analytical panel included 49 metabolites, comprising amino acids, organic acids, TCA-cycle intermediates, ketone-body-related metabolites, free carnitine, and acylcarnitine species.

After adding known amounts of labeled standards (from MSK-A2-S Metabolomics Amino Acid Mix standard, MSK-OA-1 Labeled Organic Acid Mix, and NSK-B-1 Labeled Carnitine, CIL Cambridge, MA, USA), liver samples were homogenized in 300 µL of methanol using a Precellys Evolution Homogenizer with 3 cycles of 30 s at 5800 rpm, with 20 s pauses between cycles, at 4°C (Bertin Instruments). Isotope-labeled internal standards were added before extraction to allow relative quantification and to monitor extraction efficiency. Proteins were precipitated by centrifugation for 20 min at 12,900 rpm at 4°C, and metabolites were extracted using a modified Folch method by adding 600 µL of chloroform (chloroform:methanol solution, 2:1, v/v) and 200 µL of $H_2O$, followed by centrifugation for 15 min at 12,900 rpm at 4°C. The upper aqueous phases were collected and transferred to a tube. A second extraction of the down phase was performed by adding 300 µL of methanol and 200 µL of $H_2O$ and the procedure was repeated to ensure full recovery. The total extracted amount was then dried under a gentle nitrogen stream and resuspended in 150 µL of $H_2O$ MilliQ and 50 µL of acetonitrile. Samples were first analyzed by liquid chromatography/quadrupole time-of-flight mass spectrometry in positive electrospray ionization (UHPLC-QTOF, 1290 Infinity-6545 Agilent Technology, Santa Clara, CA, USA) equipped with an Acquity BEH C18 2.1 × 100 mm 1.7-Micron column (Waters, Milford, MA, USA) for the measurement of acylcarnitines (quantified using $^{2}H$ labeled carnitine internal standards CIL Cambridge, MA, USA)

The samples were then dried again under a gentle nitrogen stream and derivatized with 10 µL of methoxyamine hydrochloride solution in pyridine (20 mg/mL; Merck KGaA) for 30 min at 60°C, followed by 30 µL of N-(tert-butyldimethylsilyl)-N-methyltrifluoroacetamide (TBDMS; Merck KGaA) and 70 µL of acetonitrile for 1 h at 60°C for amino acid and organic acid quantification by gas chromatography–tandem mass spectrometry using a GC 8890/MS 7000D system (Agilent Technologies) equipped with a DB-5MS J&W capillary column, 30 m length, 0.25 mm internal diameter, and 0.25 µm film thickness (J&W, Agilent).

The method for the identification and quantification of polar metabolites was set up based on retention-time matching against authenticated reference standards analyzed under identical chromatographic conditions and mass spectral matching against the NIST mass spectral library. Labelled internal standards used during the analysis allowed an accurate quantification and normalization across samples. Only metabolites fulfilling predefined identification and reproducibility criteria were retained for downstream analysis.

Quality control was performed using QC samples previously prepared, supplemented with the same internal standards as the experimental samples, and validated against NIST reference standards. These QC samples were injected at regular intervals throughout the analytical run to monitor instrument stability and reproducibility. The variability of the peak values of the labelled internal standard was also utilized to monitor and evaluate the overall quality of the analytical run. Metabolite labelled standards with a coefficient of variation (CV) greater than 20% across samples of the same batch were excluded from downstream analyses.

All samples were processed in two consecutive analytical batches to minimize inter-batch variability. Quantification was performed by normalizing the metabolite peak area to the corresponding internal standard peak area, followed by multiplication by the concentration of the internal standard. The resulting data were subsequently normalized to liver weight and to QC values to adjust for any minor analytical drift. A complete metabolite table, including individual values, group means, fold changes, nominal p-values, and FDR-adjusted p-values, is provided in Supplementary Table S2.

Data analysis was performed using MetaboAnalyst 6.0 software (http://www.metaboanalyst.ca/).

**Immunohistochemical analysis**

Formalin-fixed, paraffin-embedded liver sections (4 µm) were stained with hematoxylin and eosin (H&E) or processed for Ki67 immunohistochemistry. H&E staining was performed using standard protocols (05-06002/L and #05-10002/L, BioOptica). For immunohistochemistry, sections were stained with an anti-Ki67 antibody (#12202, Cell Signalling) using a Leica BOND RX automated system with heat-induced antigen retrieval (ER1 citrate buffer) and detected with Bond Polymer Refine Detection (Leica, DS9800). Slides were digitized at 20X magnification with an Aperio AT2 scanner. Ki67-positive cells were quantified using QuPath (v0.6.0) and expressed as the percentage of positive nuclei. For H&E analysis, three sections per mouse were analyzed, with three 500 × 500 µm regions of interest (ROIs) per section. Nuclei and hepatocytes were automatically detected in QuPath, with parameters optimized for nuclear size, inter-nuclear distance, and hepatocyte radius. An automated script calculated nuclei and hepatocyte densities per $mm^2$. Outputs were quality-controlled to ensure consistent and reliable quantification of hepatocyte and nuclear density across experimental conditions.

**Glucose and Insulin Tolerance Analysis**

Glucose (GTT) and insulin (ITT) tolerance tests were performed in NP and GD15 females after overnight (GTT) or 4h (ITT) fasting. Mice received intraperitoneal injections of glucose (2 g/kg) or insulin (0.75 IU/kg), and blood glucose levels were measured at baseline and 15, 30, 60, 90, and 120 min post-injection using a glucose meter (Accu-Chek® Instant Meter, Roche).

**Western Blotting Analysis**

Frozen mouse livers were homogenized in ice-cold buffer (20 mM HEPES, 5 mM $MgCl_2$, 420 mM NaCl, 0.1 mM EDTA, and 20% glycerol) containing protease and phosphatase inhibitors (Pierce™ Protease Inhibitor Mini Tablets, EDTA-free, A32955, Thermo Scientific™ Waltham). Homogenates were centrifuged at 12000 g for 20 min at 4 °C, and the supernatants were collected. After protein quantification (Pierce™ Bradford Protein Assay Kit, 23200, Thermo Scientific™ Waltham), 25 µg of proteins were resolved on 10% SDS–PAGE gels and transferred to nitrocellulose membranes (Amersham Protran 0.45 NC, 10600018). Membranes were incubated overnight at 4 °C with primary antibodies followed by HRP-conjugated secondary antibodies for 1 h at room temperature (RT). Primary antibodies included anti-ERα (Abcam, AB32063); anti-β-actin (Sigma Aldrich, Merck KGaA, A1978); anti-p-AKT Ser473 (#4060), anti-AKT ( #4691), anti-p-GSK3β Ser9 (#5558), anti-GSK3β (#12456), anti-p-mTOR Ser2448 (#5536), anti-mTOR (#2983), anti-p-S6K1 Thr389 (#9234), anti-S6K1 (#2708), anti-p-S6 Ser235/236 (#4858), anti-S6 (#2217), all from Cell Signaling. Signals were detected using enhanced chemiluminescence (Cytiva Italy SRL, RPN2209) and acquired with an Odyssey Fc Imaging System; band intensities were quantified using Image Studio software (LiCor Biosciences).

**Metabolic cages and indirect calorimetry**

Indirect calorimetry was performed using a computer-controlled Promethion Metabolic Screening system (Sable Systems International). Mice were singly housed in metabolic cages and acclimatized for 48 h before data acquisition. Additional analyses comparing 48-, 72-, and 84-h acclimatization periods did not reveal major differences in the metabolic parameters measured. Food and water were available ad libitum throughout the study. Oxygen consumption ($VO_2$), carbon dioxide production ($VCO_2$), respiratory exchange ratio (RER), locomotor activity, food intake, and energy expenditure were continuously monitored over a 48-h recording period under a 12-h light/dark cycle. Respiratory gases were measured using integrated $O_2$, $CO_2$, and water vapor analyzers (GA3m1, Sable Systems International), with measurements acquired for 1 min every 7 min. Data were analyzed using the web tool Indirect Calorimetry Experiments CalR and represented as hourly averages and as light- and dark-phase summaries. Raw data and analyzed outputs from indirect calorimetry are provided in the Supplementary Table S3.

### Statistical analysis

Statistical analyses were performed using GraphPad Prism 8.0. Data are presented as mean ± SEM. Exact group-specific sample sizes are reported in the figure legends and summarized in Supplementary Table S4; each biological replicate represents an individual animal unless otherwise specified. Two-group comparisons were analyzed using two-tailed unpaired Student's t-tests, whereas multiple-group datasets were analyzed using one- or two-way ANOVA followed by Bonferroni-adjusted *post hoc* comparisons. For factorial experimental designs, two-way ANOVA was used to assess the main effects of the two factors relevant to each experiment and their interaction. For the acute insulin-stimulation experiment, genotype × treatment effects were analyzed separately within each reproductive stage. Longitudinal GTT and ITT glucose profiles were analyzed using mixed-design repeated-measures ANOVA, with experimental group as the between-subject factor and time as the within-subject factor; AUC values were analyzed separately by two-way ANOVA. Phase-averaged indirect calorimetry data were analyzed using a mixed-design repeated-measures ANOVA, with experimental group as the between-subject factor and light/dark phase as the within-subject factor, followed by Bonferroni-adjusted prespecified comparisons.

For RNA-seq analyses, differentially expressed genes were identified using FDR-adjusted P-values, as described in the Transcriptomics data analysis section. For metabolomic analyses, nominal and FDR-adjusted P-values are reported in the corresponding supplementary tables.

All tests were two-sided, with $p < 0.05$ considered statistically significant.

**Supplementary figure S1**

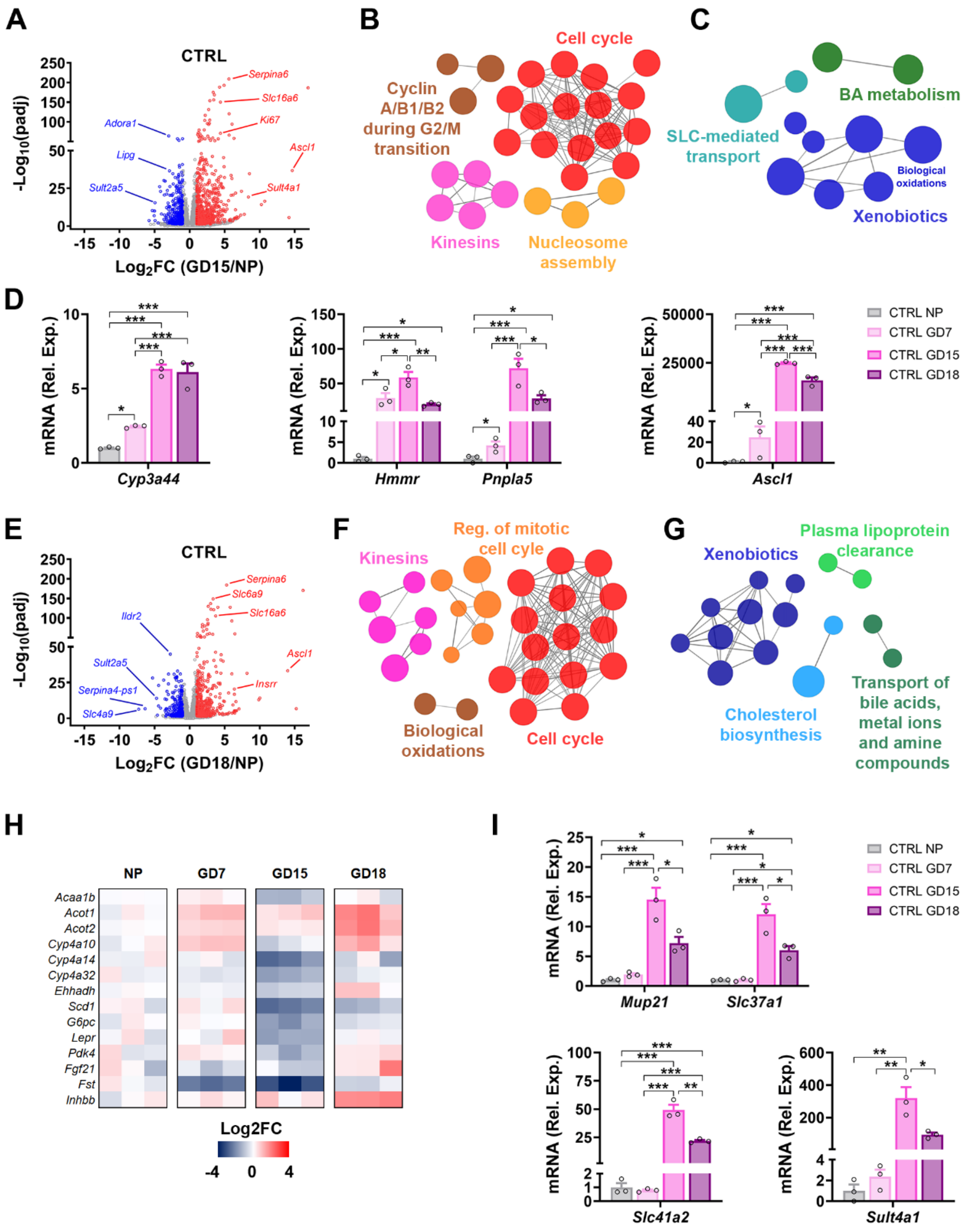


**Fig. S1. Stage-specific transcriptomic remodeling in CTRL liver during pregnancy**. (**A**) Volcano plot of liver DEGs measured at mid-pregnancy, comparing GD15 with NP CTRL. (**B-C**) GO analysis of the functional networks significantly up- (**B**) and down-regulated (**C**) in CTRL liver at GD15 compared with NP. (**D**) mRNA levels of *Cyp3a44*, *Hmmr*, *Pnpla5*, and *Ascl1* measured in CTRL liver during pregnancy. (**E**) Volcano plot of liver DEGs at late pregnancy, comparing GD18 with NP CTRL. (**F-G**) GO analysis of the functional networks significantly up- (**F**) and down-regulated (**G**)

in CTRL liver at GD18 compared with NP. (**H**) Heatmap showing PPARα target genes with reduced expression at GD15 and re-induction at GD18 in CTRL liver. (**I**) mRNA levels of *Mup21*, *Slc37a1*, *Slc41a2*, and *Sult4a1* measured in CTRL liver during pregnancy.
Normalized RNA-seq expression values in Fig. S1D and S1I are shown as mean ± SEM (n = 3). *p<0.05, **p<0.01 and ***p<0.001 by one-way ANOVA followed by Bonferroni's *post hoc* test.

### Supplementary figure S2

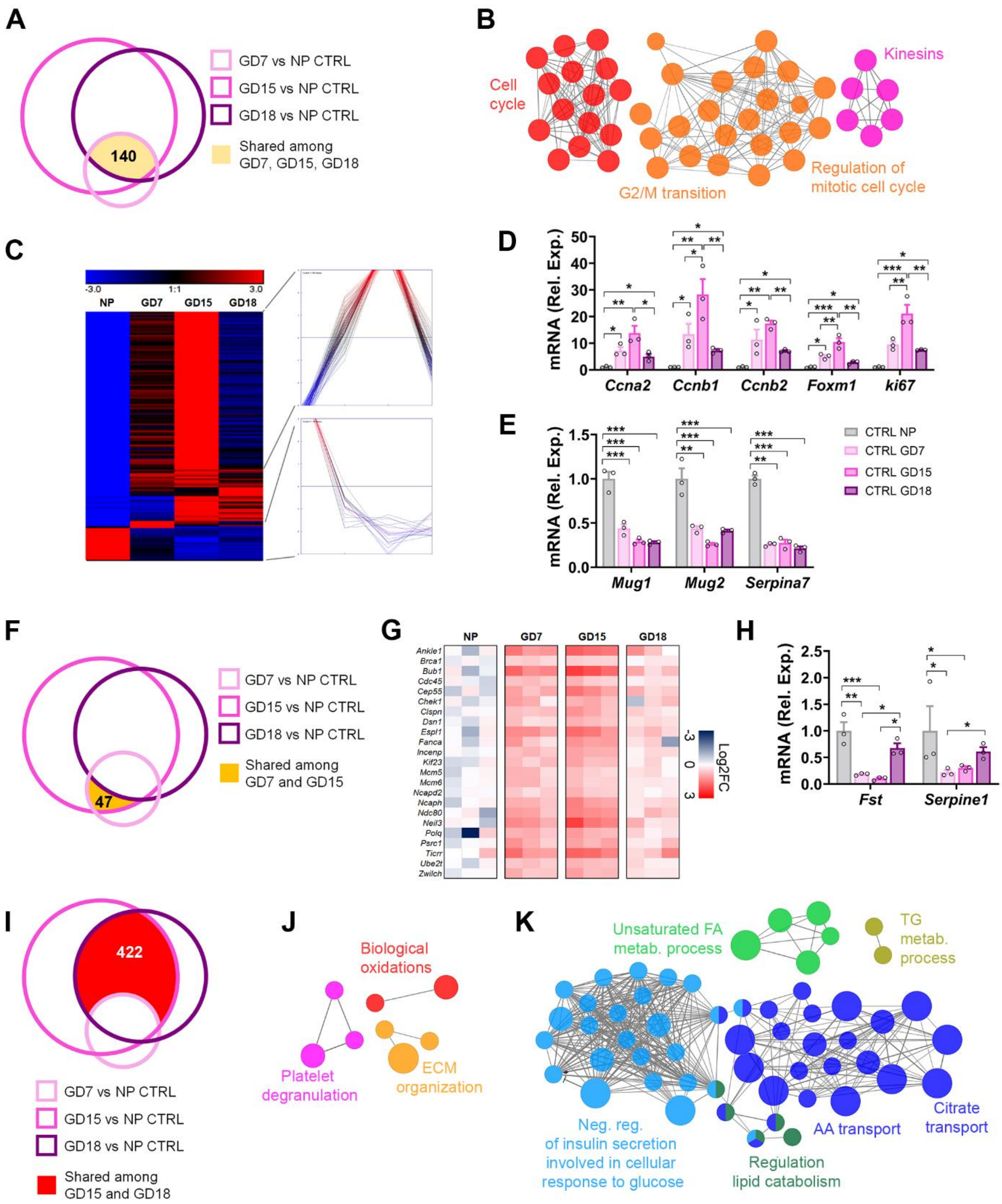


**Fig. S2. Shared and stage-specific liver transcriptomic signatures during pregnancy**. (**A**) Venn diagram showing shared liver DEGs across GD7, GD15 and GD18 CTRL compared with NP CTRL. (**B**) GO analysis of the functional networks significantly up-regulated in the liver at GD7, GD15 and GD18 compared with NP CTRL. (**C**) Heatmap showing shared liver DEGs among GD7, GD15 and GD18 CTRL compared with NP CTRL. (**D-E**) Representative liver genes up- (**D**) and down-regulated

(**E**) at all pregnancy stages compared with NP. (**F**) Venn diagram showing shared liver DEGs between GD7 and GD15 compared with NP CTRL. (**G**) Heatmap showing the genes up-regulated at GD7 and GD15 compared with NP CTRL. (**H**) mRNA levels of *Fst* and *Serpine1* in CTRL liver during pregnancy. (**I**) Venn diagram showing shared liver DEGs between GD15 and GD18 compared with NP CTRL. (**J, K**) GO analysis of the functional networks significantly up- (**J**) and down- (**K**) regulated in the liver at GD15 and GD18 compared with NP CTRL. Normalized RNA-seq expression values in Fig. S2D, S2E, and S2H are shown as mean ± SEM (n = 3). *$p<0.05$, **$p<0.01$ and ***$p<0.001$ by one-way ANOVA followed by Bonferroni's *post hoc* test.

### Supplementary figure S3

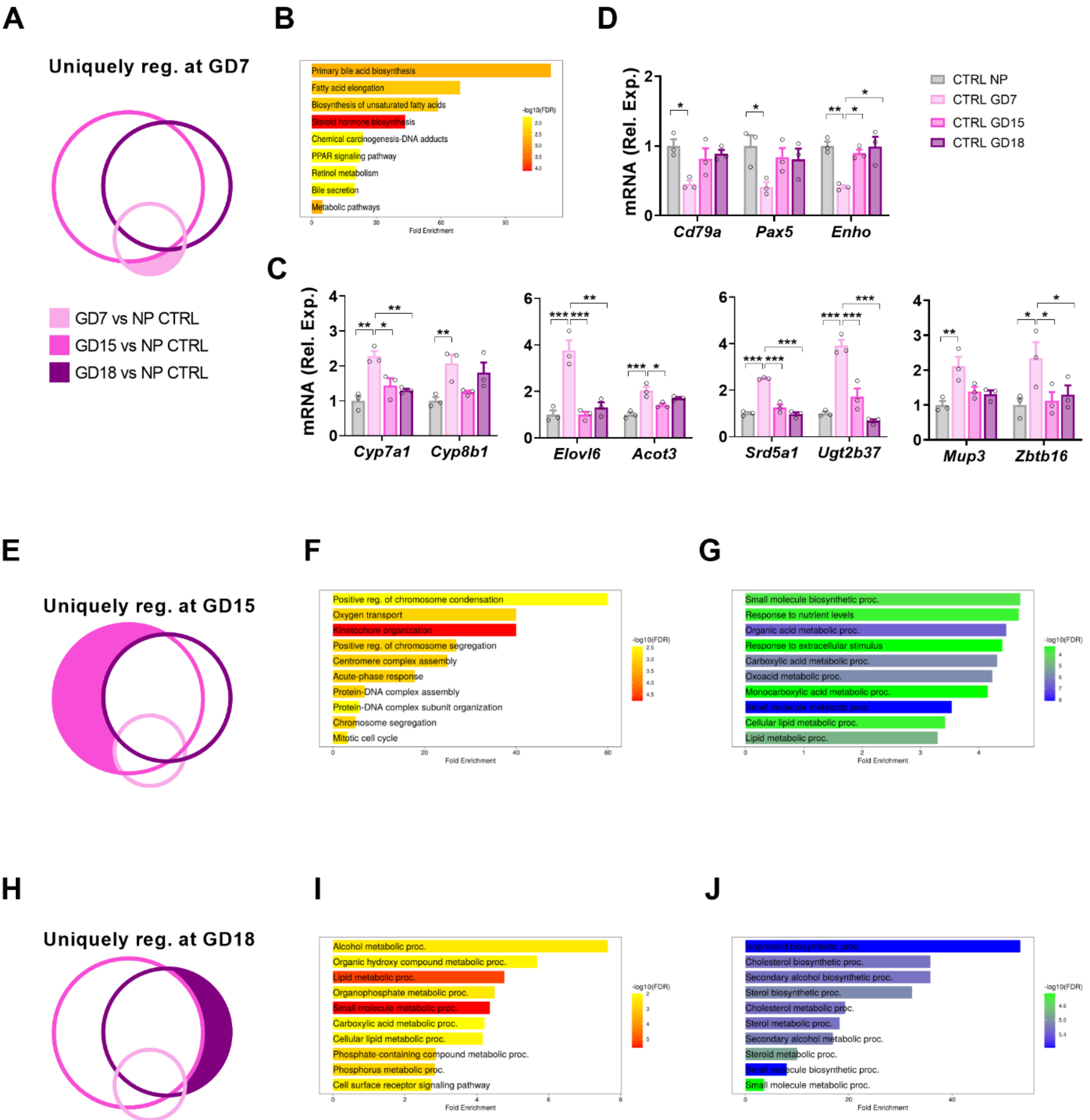


**Fig. S3. Gestational stage-specific regulation of the CTRL liver transcriptome**. (**A**) Venn diagram highlighting in pink the subset of liver genes uniquely regulated in early pregnancy (GD7). (**B**) GO analysis of the functional networks up-regulated uniquely in CTRL liver at GD7 compared with NP. (**C**) mRNA levels of genes involved in bile acid (BA), fatty acid (FA), steroid, and glucose metabolism specifically induced in CTRL liver at GD7. (**D**) mRNA levels of immune-related genes and *Enho* specifically reduced in CTRL liver at GD7. (**E**) Venn diagram highlighting in fuchsia the subset of liver genes uniquely regulated in mid-pregnancy (GD15). (**F-G**) GO analysis of the functional networks up- (**F**) and down-regulated (**G**) in CTRL liver uniquely at GD15 compared with

NP. (**H**) Venn diagram highlighting in violet the subset of liver genes uniquely regulated in late pregnancy (GD18). (**I-J**) GO analysis of the functional networks up- (**I**) and down-regulated (**J**) uniquely in CTRL liver at GD18 compared with NP. Normalized RNA-seq expression values in Fig. S3C and S3D are shown as mean ± SEM (n = 3). *$p<0.05$, **$p<0.01$ and ***$p<0.001$ by one-way ANOVA followed by Bonferroni's *post hoc* test.

### Supplementary figure S4

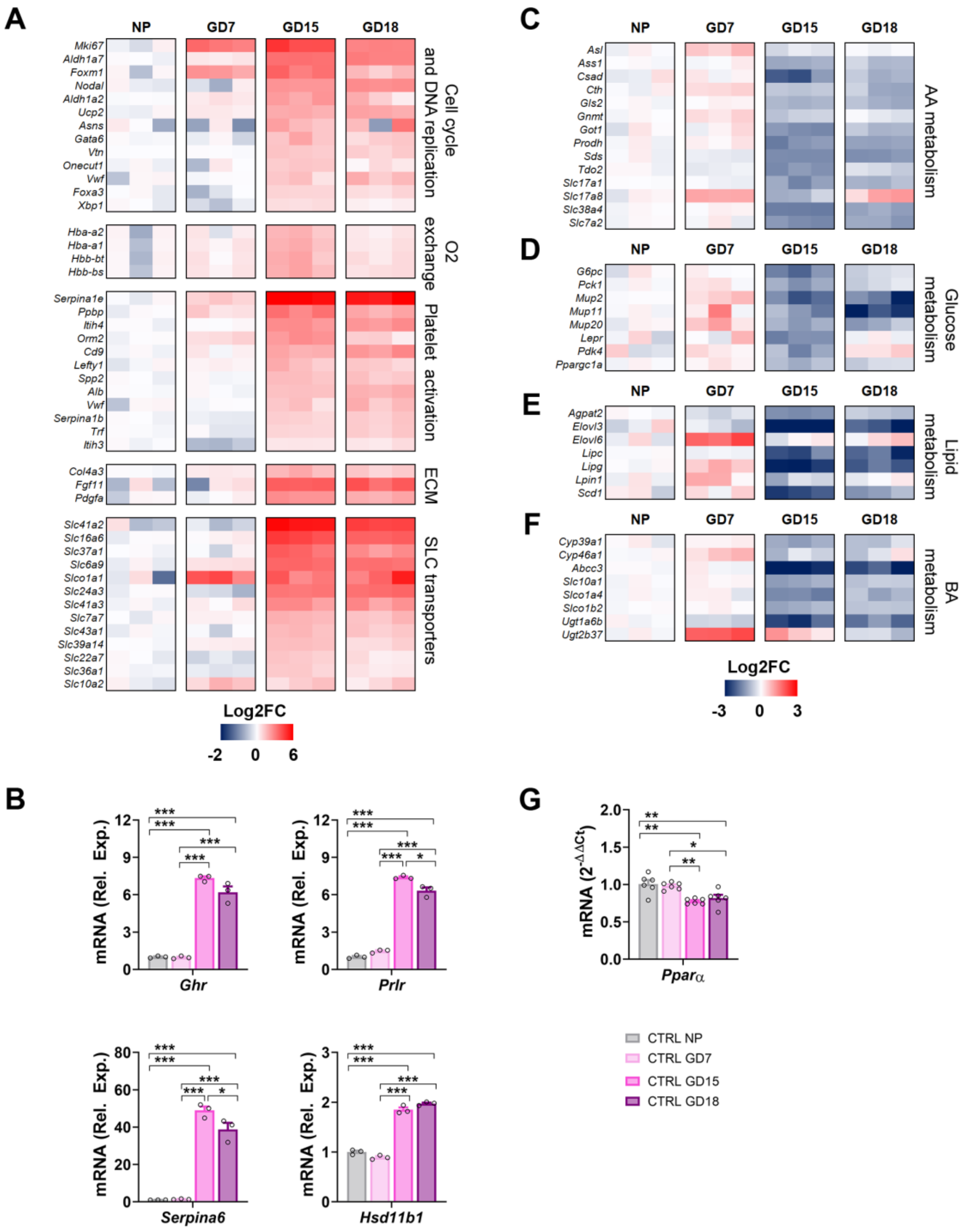


**Fig. S4. Mid-gestation is associated with proliferation-related gene expression and metabolic remodeling in CTRL liver**. (**A**) Heatmap showing genes involved in cell-cycle and DNA replication, oxygen transport, platelet activation, extracellular matrix (ECM) remodeling and angiogenesis, and transport (SLC proteins) induced in CTRL liver in mid- and late pregnancy. (**B**) mRNA levels of *Ghr*, *Prlr*, *Serpina6*, and *Hsd11b1* induced in CTRL liver in mid and late pregnancy. (**C-F**) Heatmaps showing genes involved in amino acid (**C**), glucose (**D**), lipid (**E**), and bile acid (**F**) metabolism with reduced expression in CTRL liver at mid-pregnancy. (**G**) mRNA levels of *Pparα* in CTRL liver during pregnancy. Data in Fig. S4B and S4G are shown as mean ± SEM (n=3). **p<0.01 and ***p<0.001 by one-way ANOVA followed by Bonferroni's *post hoc* test.

### Supplementary figure S5

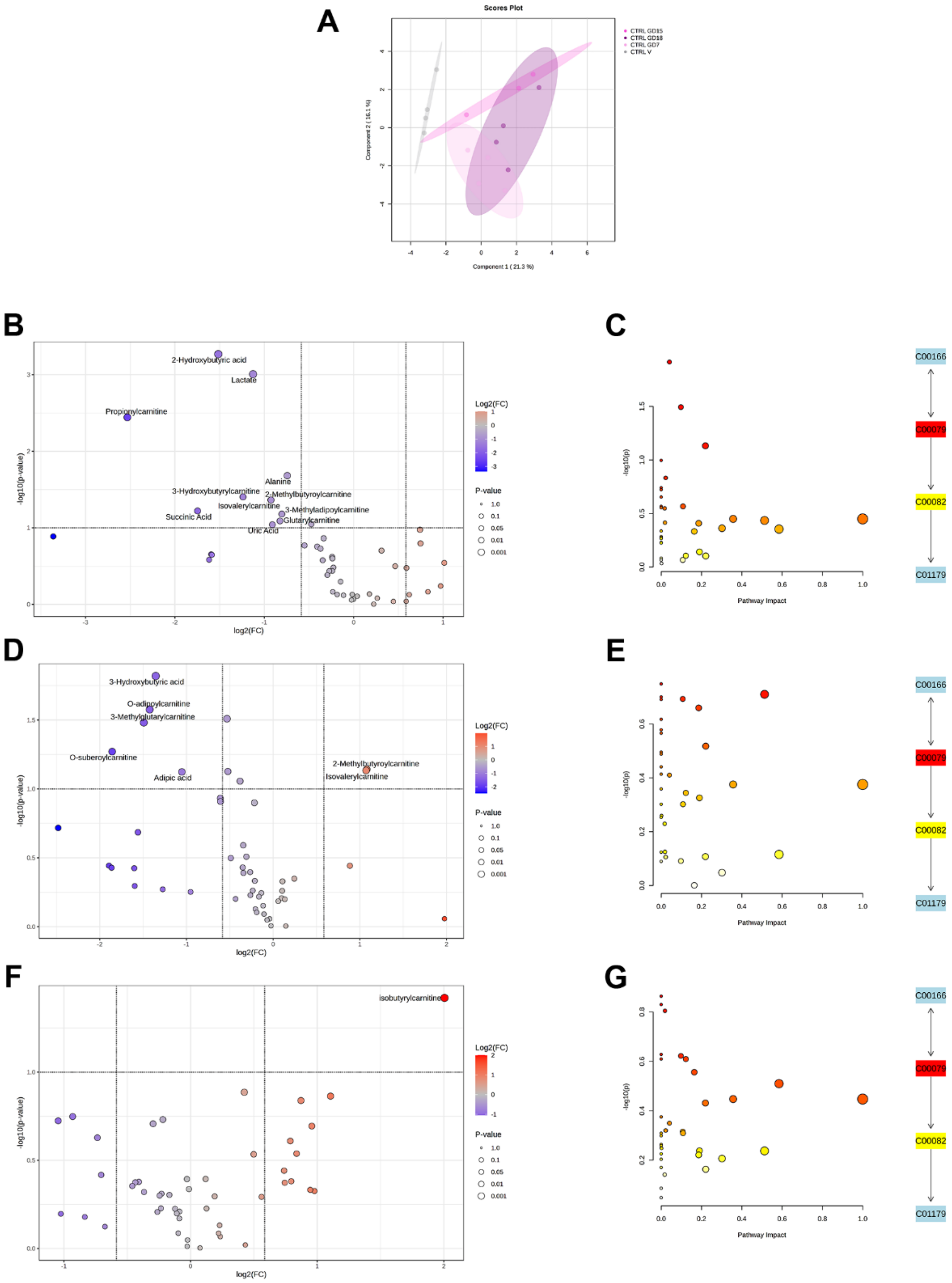


**Fig. S5. Targeted metabolomic changes in CTRL liver across pregnancy transitions.** (**A**) Principal component analysis (PCA) of targeted metabolites measured in CTRL liver during pregnancy. (**B-C**) Volcano plot (**B**) and pathway impact analysis (**C**) of targeted metabolites measured in NP and GD7 CTRL livers. (**D-E**) Volcano plot (**D**) and pathway impact analysis (**E**) of targeted metabolites measured in GD7 and GD15 CTRL livers. (**F-G**) Volcano plot (**F**) and pathway impact analysis (**G**) of targeted metabolites measured in GD15 and GD18 CTRL livers. Targeted metabolomic analysis was performed with n = 4 biological replicates per group. In volcano plots, significantly increased and decreased metabolites are shown in red and blue, respectively; other metabolites are shown in gray. In pathway impact plots, metabolic pathways are represented as circles according to enrichment p-value and pathway impact using MetaboAnalyst 6.0. Circle color reflects the p-value from enrichment analysis, and circle size represents the fold enrichment score.

**Supplementary figure S6**

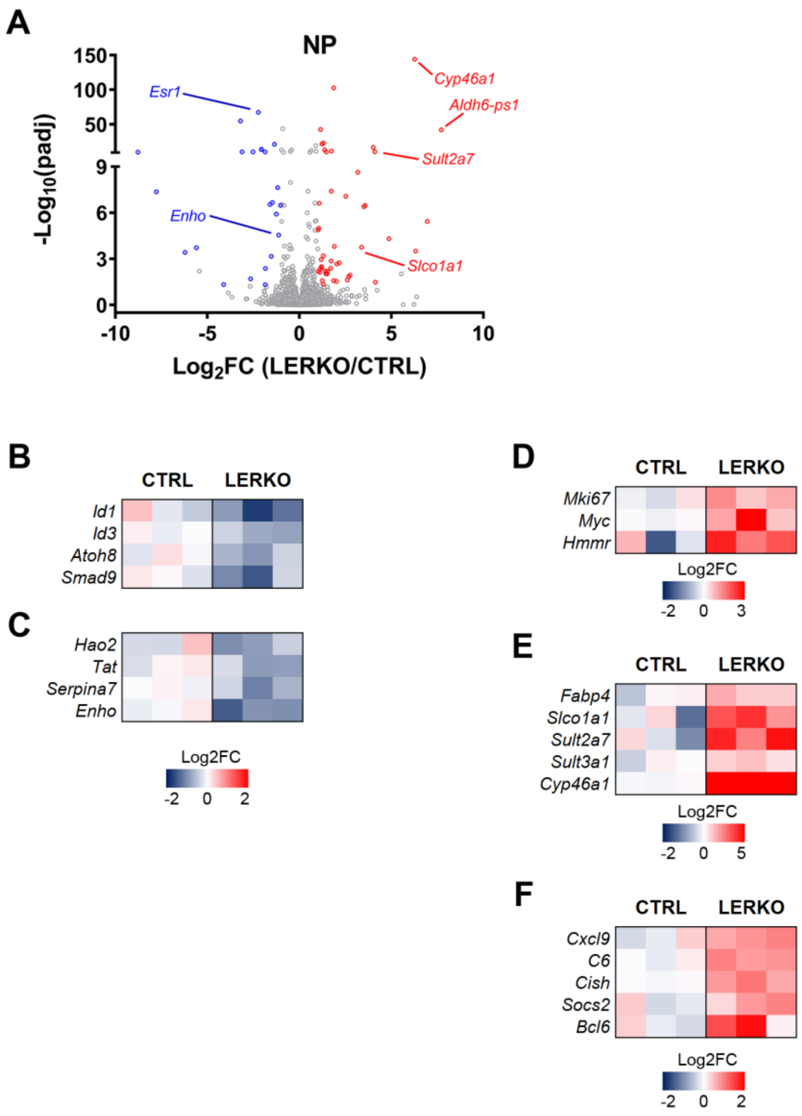


**Fig. S6. Baseline transcriptional differences between CTRL and LERKO livers under non-pregnant conditions. (A)** Volcano plot showing DEGs between CTRL and LERKO NP livers. **(B-C)** Heatmaps of selected differentially expressed genes showing reduced expression of estrogen-responsive and hepatocyte differentiation/quiescence-associated genes (**B**), and genes involved in lipid and energy metabolism and hepatokine signaling (**C**) in LERKO livers compared with CTRL. (**D-F**) Heatmaps showing increased expression of genes associated with cell-cycle progression and proliferative activity (**D**), lipid and sterol transport and metabolism (**E**), and immune or cytokine-responsive pathways (**F**) in LERKO livers compared with CTRL. Normalized RNA-seq expression values are shown as means (n = 3).

### Supplementary figure S7

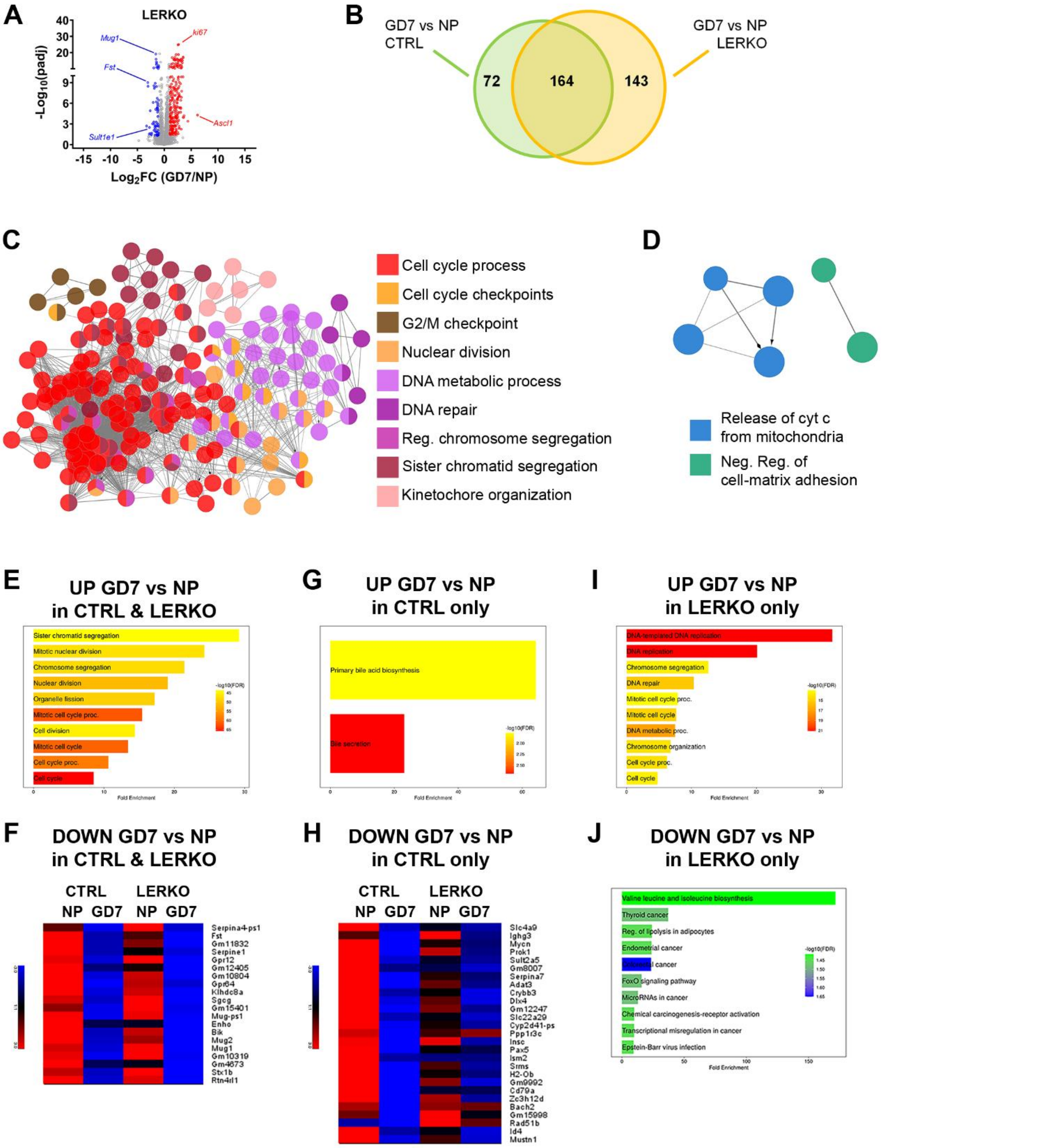


**Fig. S7. ERα deficiency alters early pregnancy-associated liver transcriptomic adaptation.** (**A**) Volcano plot of DEGs measured in LERKO liver during early pregnancy (GD7). (**B**) Venn diagram showing the overlap between liver DEGs in CTRL and LERKO mice in the NP-to-GD7 transition. (**C-D**) GO analysis of the functional networks up- (**C**) and down- (**D**) regulated in LERKO liver at

GD7 compared with NP LERKO. (**E**) GO analysis of the functional networks up-regulated in both CTRL and LERKO livers at GD7 compared with their NP counterparts. (**F**) Heatmap showing genes down-regulated in both CTRL and LERKO livers at GD7 compared with their NP counterparts. (**G**) GO analysis of the functional networks up-regulated uniquely in CTRL liver at GD7 compared with NP CTRL. (**H**) Heatmap representing the genes down-regulated uniquely in CTRL liver at GD7 compared with NP CTRL. (**I-J**) GO analysis of the functional networks up- (**I**) and down- (**J**) regulated uniquely in LERKO liver at GD7 compared with their NP LERKO. Normalized RNA-seq expression values are shown as means (n = 3).

### Supplementary figure S8

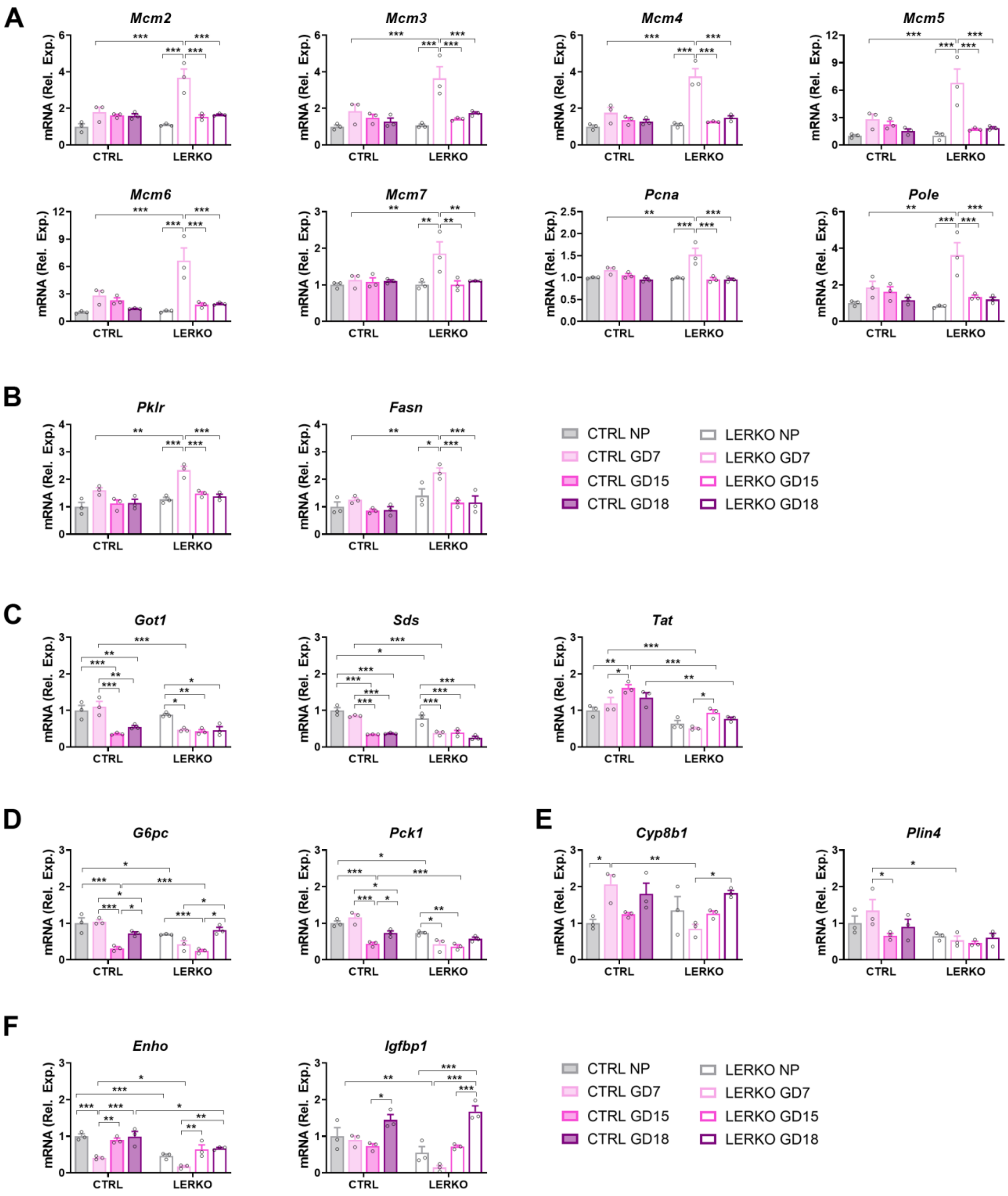


**Fig. S8. Early pregnancy in LERKO livers is associated with altered proliferative and metabolic gene expression.** (**A-F**) mRNA levels of genes differentially regulated in LERKO liver compared with CTRL at GD7, involved in cell cycle, DNA replication and repair (**A**), glycolysis and *de novo* lipogenesis (**B**), amino acid catabolism (**C**), gluconeogenesis (**D**), PPARα signaling (**E**), and hepatokine signaling (**F**). Data are shown as mean ± SEM (n=3). *p<0.05, **p<0.01 and ***p<0.001 by two-way ANOVA followed by Bonferroni's *post hoc* test.

### Supplementary figure S9

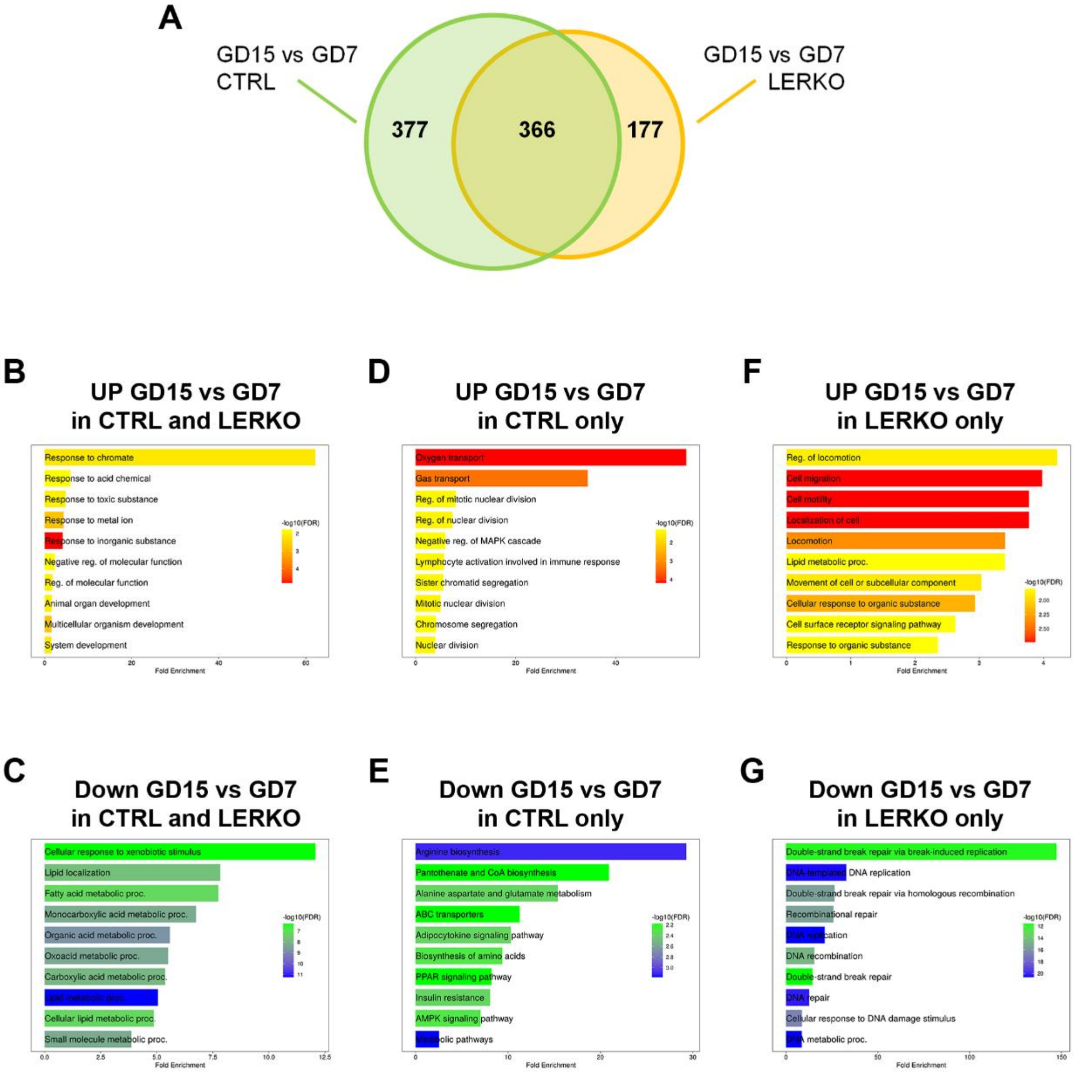


**Fig. S9. Hepatic ERα deficiency is associated with altered proliferative transcriptional responses at mid-gestation.** (**A**) Venn diagram showing the overlap of liver DEGs in CTRL and LERKO females during the GD7-to-GD15 transition. (**B-C**) GO analysis of the functional networks up- (**B**) and down-regulated (**C**) in the liver of both CTRL and LERKO females during the GD7-to-GD15 transition. (**D-E**) GO analysis of the functional networks up- (**D**) and down-regulated (**E**) uniquely in GD15 CTRL liver compared with GD7 CTRL. (**F-G**) GO analysis of the functional networks uniquely up- (**F**) and down-regulated (**G**) in GD15 LERKO liver compared with GD7 LERKO.

**Supplementary figure S10**

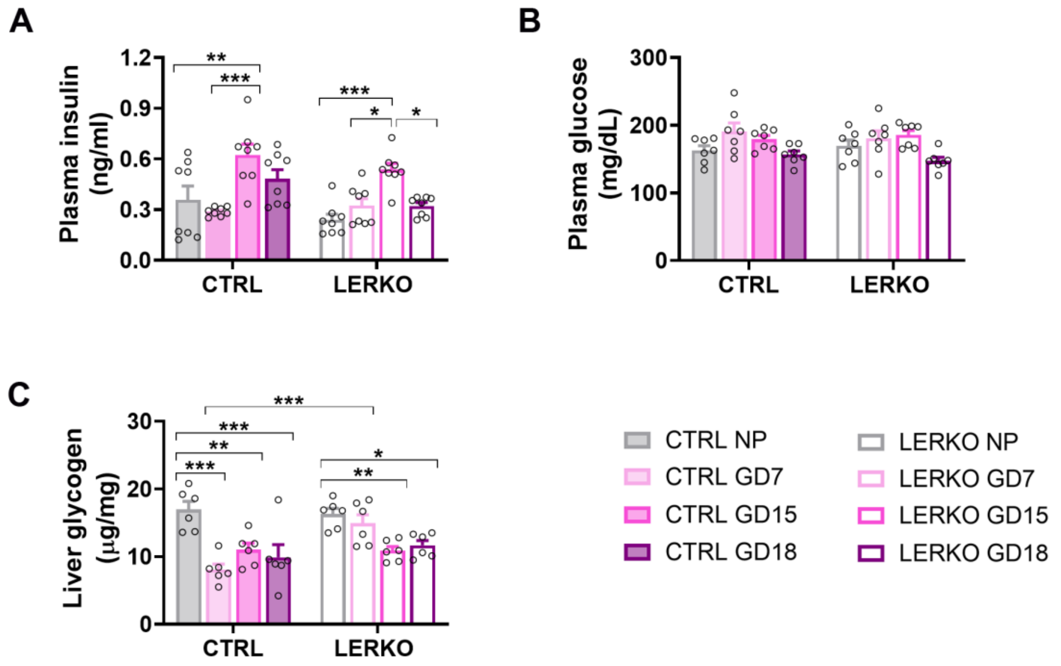


**Fig. S10. Hepatic ERα deficiency is associated with altered hepatic glycogen content during early pregnancy.** (**A-B**) Plasma insulin (**A**, n=8) and glucose (**B**, n=7) levels measured in NP and pregnant CTRL and LERKO at GD7, GD15 and GD18. (**C**) Glycogen content measured in CTRL and LERKO livers during pregnancy (n=6). Data are mean ± SEM; *p<0.05, **p<0.01 and ***p<0.001 by two-way ANOVA followed by Bonferroni's *post hoc* test.

**Supplementary figure S11**

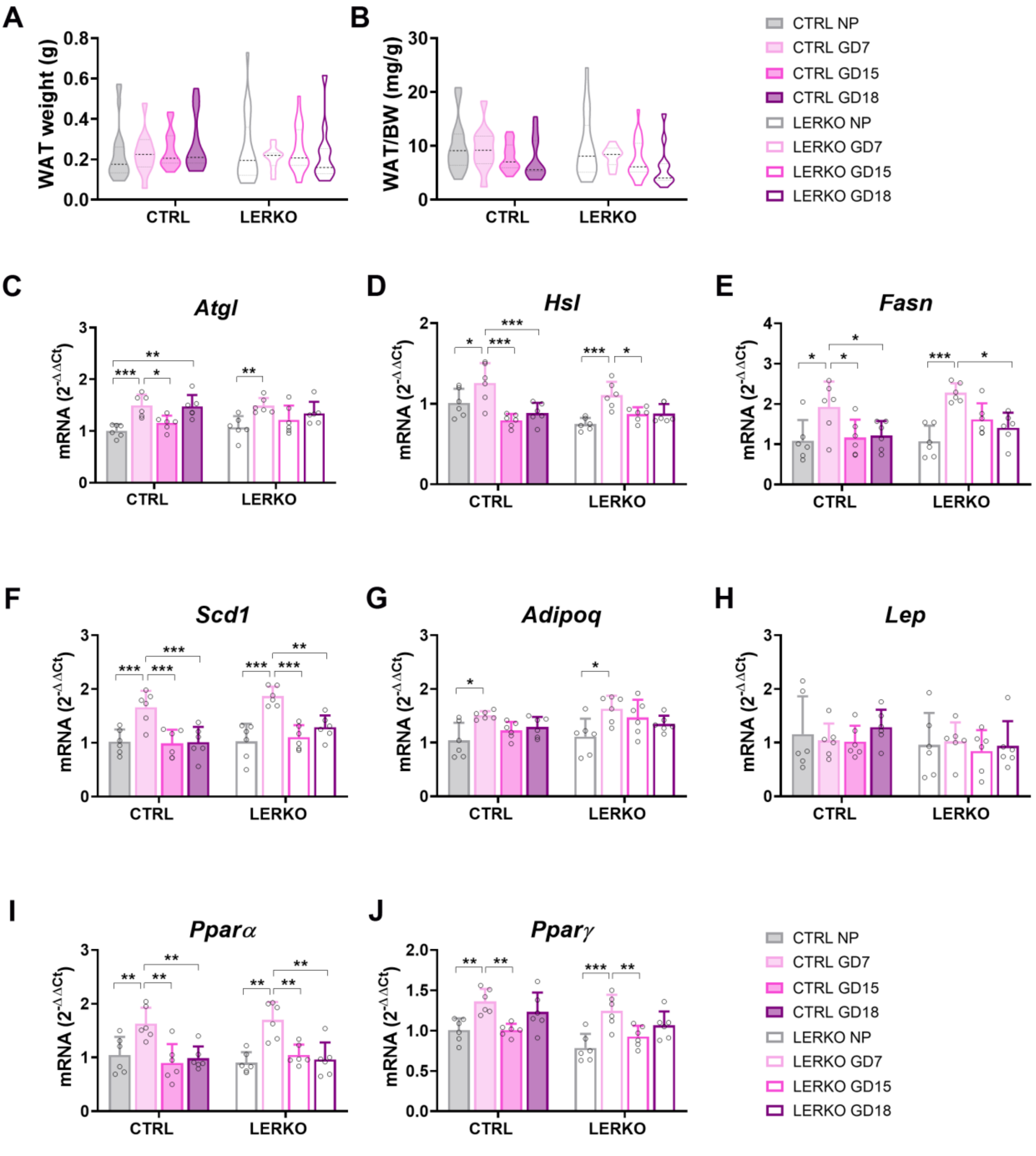


**Fig. S11. Visceral WAT mass and expression of selected adipose metabolic regulators are largely preserved in LERKO females.** (**A-B**) Visceral WAT weight expressed in grams (**A**) or normalized to BW (**B**) in NP and pregnant CTRL and LERKO mice (CTRL NP/GD7/GD15/GD18, n=20/16/18/16, and LERKO NP/GD7/GD15/GD18, n=20/16/18/16). (**C-J**) mRNA levels of *Atgl*, *Hsl*, *Fasn*, *Scd1*, *Adipoq*, *Lep*, *Pparα* and *Pparγ* measured by qPCR in CTRL and LERKO WAT during pregnancy (n=6). Data are mean ± SEM; *p<0.05, **p<0.01 and ***p<0.001 by two-way ANOVA followed by Bonferroni's post hoc test.

### Supplementary figure S12

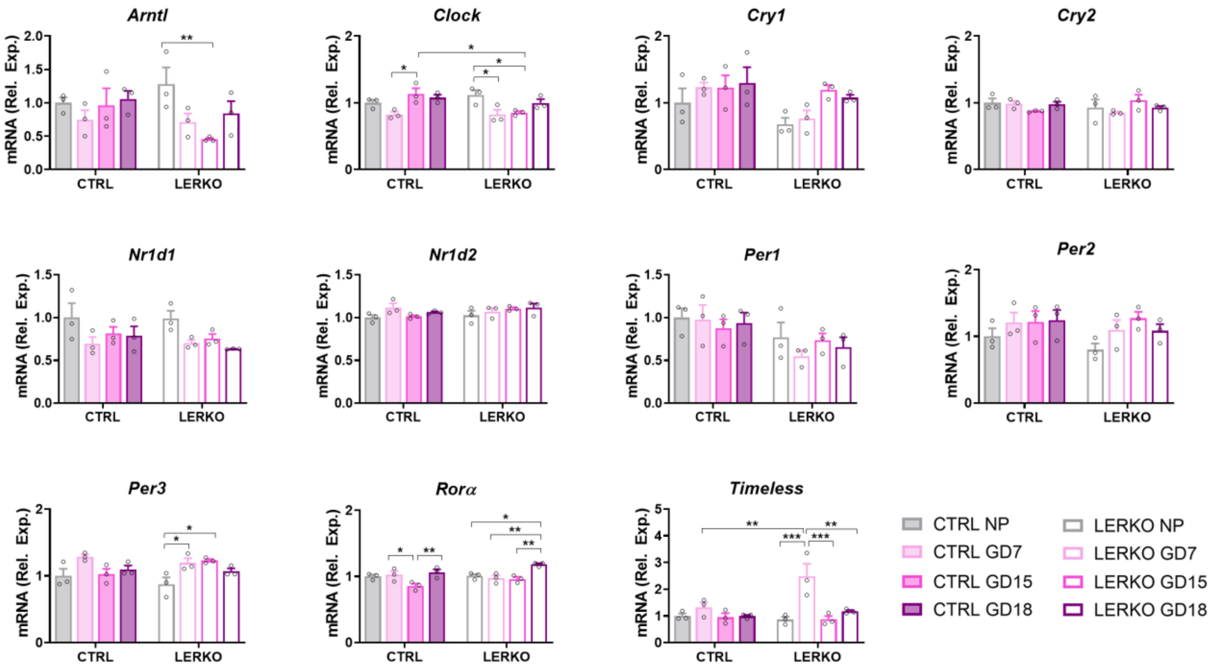


**Fig. S12. Hepatic ERα deficiency is associated with altered expression of selected liver clock genes at the analyzed time point.** Normalized RNA-seq expression levels of selected clock genes measured in CTRL and LERKO livers during pregnancy. Data are shown as mean ± SEM (n = 3). *p<0.05, **p<0.01 and ***p<0.001 by two-way ANOVA followed by Bonferroni's *post hoc* test.
